# Identifying strengths and weaknesses of methods for computational network inference from single cell RNA-seq data

**DOI:** 10.1101/2021.06.01.446671

**Authors:** Sunnie Grace McCalla, Alireza Fotuhi Siahpirani, Jiaxin Li, Saptarshi Pyne, Matthew Stone, Viswesh Periyasamy, Junha Shin, Sushmita Roy

## Abstract

Single-cell RNA-sequencing (scRNA-seq) offers unparalleled insight into the transcriptional programs of different cellular states by measuring the transcriptome of thousands of individual cells. An emerging problem in the analysis of scRNA-seq is the inference of transcriptional gene regulatory networks and a number of methods with different learning frameworks have been developed to address this problem. Here, we present an expanded benchmarking study of eleven recent network inference methods on six published scRNA-seq datasets in human, mouse, and yeast considering different types of gold standard networks and evaluation metrics. We evaluate methods based on their computing requirements as well as on their ability to recover the network structure. We find that, while most methods have a modest recovery of experimentally derived interactions based on global metrics such as Area Under the Precision Recall curve, methods are able to capture targets of regulators that are relevant to the system under study. Among the top performing methods that use only expression were SCENIC, PIDC, MERLIN or Correlation. Addition of prior biological knowledge and the estimation of transcription factor activities resulted in the best overall performance with the Inferelator and MERLIN methods that use prior knowledge outperforming methods that use expression alone. We found little to no benefit from imputation for network inference, which is further dataset-dependent. Comparisons of inferred networks for comparable bulk conditions showed that the networks inferred from scRNA-seq datasets are often better or at par with the networks inferred from bulk datasets. Our analysis should be beneficial in selecting methods for network inference. At the same time, this highlights the need for improved methods and better gold standards for regulatory network inference from scRNAseq datasets.

## Introduction

Inference of genome-scale regulatory networks from global mRNA profiles has been a long-standing problem in systems biology and gene regulation. These networks are important for understanding the molecular mechanisms of diverse biological processes, including response to environmental signals, cell type specification and disease. The availability of single-cell omic data opens up new opportunities to reverse engineer transcriptional regulatory networks, enabling us to study the regulatory network from multiple cell types. Single-cell RNA-seq (scRNA-seq) measures genome-wide expression profiles of tens of thousands of cells, which greatly reduces the cost of generating datasets with the large sample sizes needed for computational network inference. Furthermore, it offers the potential to infer cell-type-specific regulatory networks for both known and novel cell populations in the same experiment, offering an efficient way to dissect heterogeneous populations.

The availability of scRNA-seq datasets has fueled the development of a number of methods for network inference from these data that use different types of models ranging from Gaussian graphical models [1], information theoretic methods [2–4], random forests [5], ordinary differential equations [6], and Boolean networks [7] (See [8,9] for comprehensive reviews). Methods also vary in their inclusion of pseudotime [3,6,10] or imputed, de-noised signals [4]. Some of these methods specifically model the statistical properties of scRNA-seq data [11,12], while others are adaptations of existing methods for bulk data [5]. In parallel with method development, there have been two benchmarking studies [13, 14] to compare existing network inference algorithms. One study showed that existing methods perform poorly on both simulated and real data [14], while the second showed that methods that perform well on simulated data can be relatively poor on real data [13]. While these studies have been useful, a number of open questions need to be addressed to gain a comprehensive understanding of the strengths and weaknesses of existing network inference algorithms from scRNA-seq datasets. In particular, existing benchmarking efforts have studied relatively small number of networks from real data (<2000 genes), while in reality genome-scale regulatory networks can have between 5-10k genes. Another unknown is the extent to which different gold standards (e.g. ChIP-chip/seq versus regulator perturbation) can affect measured performance and to what extent do methods agree with each other on novel predictions and recovery of the gold standard networks. In addition, imputation of sparse scRNA-seq count matrices has been suggested to improve downstream analysis including identification functional relationships among genes [4]. However, it is not clear whether this benefits network inference methods more generally. Finally, experiments from bulk RNA-seq datasets have shown that integrating auxiliary sources of data to inform the graph structure and estimate transcription factor activity (TFA) levels has been useful, but the utility of prior knowledge has not been assessed for scRNA-seq network inference.

To address these gaps, we first compared 11 network inference methods on six published scRNA-seq datasets from human, mouse, and yeast (**Figure 1**). Benchmarking the algorithms for time and memory consumption on datasets of different sizes identified several algorithms that are unlikely to scale to genomewide gene regulatory networks. We compare the algorithm performance using global network topology metrics such as Area Under the Precision Recall (AUPR)) curve and F-score as well as local metrics such as the number of regulators whose targets can be predicted accurately. Based on global metrics, network inference methods had modest performance compared to random, however the local metrics show that methods can predict targets of several relevant regulators. Generally, the top performing algorithms tend to agree with each other on predicted edges, although this depends upon the depth of the dataset. Imputation of scRNA-seq datasets did not benefit majority of the network inference approaches, but was beneficial for a few methods in dataset-specific manner. To our surprise, simple correlation based methods performed as well or better on these metrics compared to many network inference algorithms. We compared the performance of these algorithms on single cell datasets to those on bulk datasets for matched conditions and find comparable performance for both classes of algorithms. We also explored the effects of dataset size (i.e. the number of cells in a dataset) on network inference and observed that the sequencing depth had a greater effect on network inference when controlled for the dataset size. Finally, methods that incorporated priors and TFA had the best overall performance across different metrics, outperforming methods that use expression alone. Overall, our study presents an extended comparison of recent algorithms of network inference from scRNA-seq datasets and identifies common strengths and weaknesses on different types of gold standards. Our results should be beneficial to the community for selecting an algorithm for network inference as well as for developing new network inference algorithms for scRNA-seq datasets.

**Figure 1.**
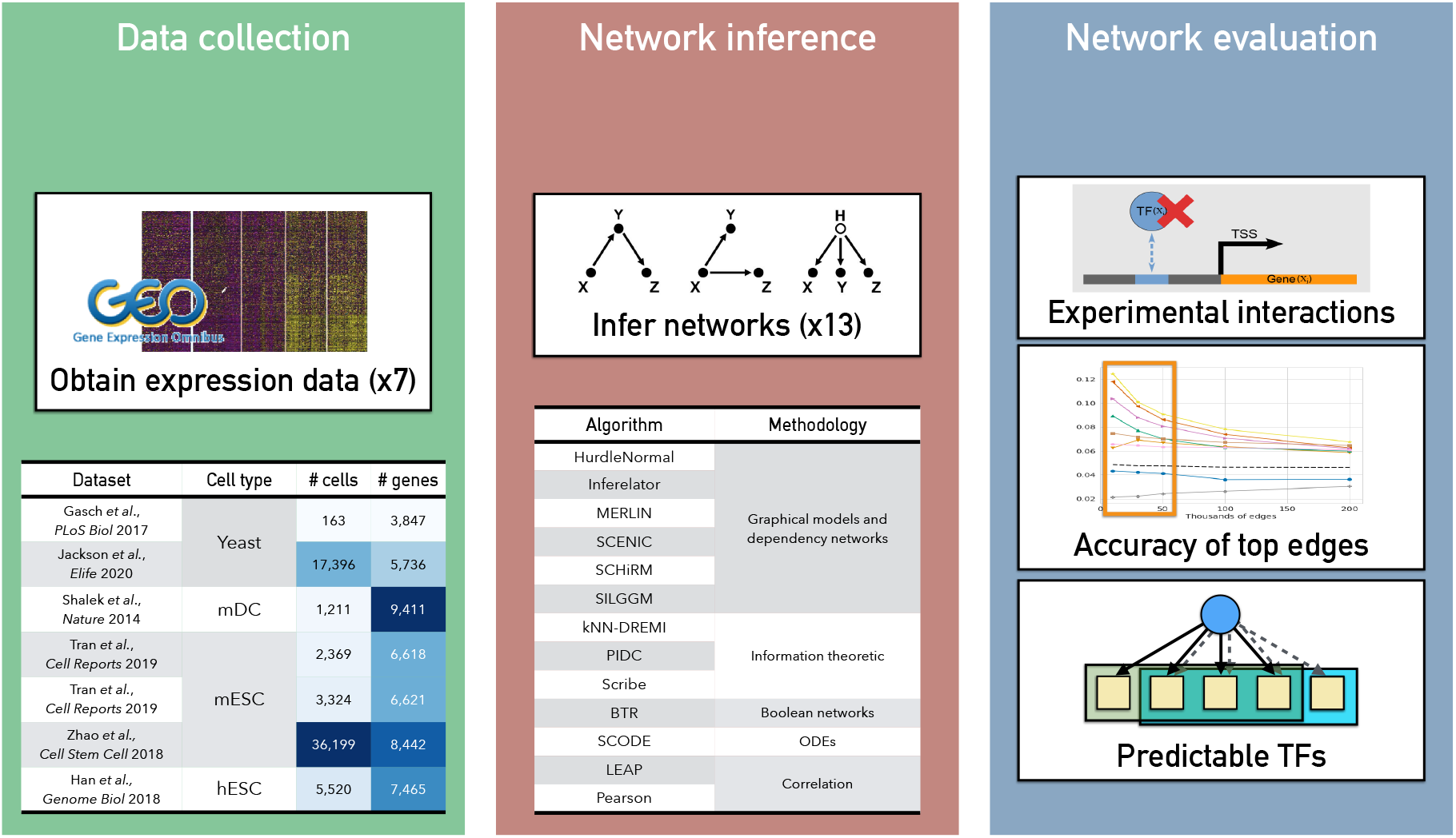
Overview of network inference benchmarking study. Our study examined 11 algorithms developed for scRNA-seq datasets and 2 algorithms for bulk expression and across 7 different scRNA-seq datasets from three species: yeast, mouse and human.

## Results

### Computing requirements of regulatory network inference algorithms for single cell RNA-seq data

We started with 11 algorithms specifically designed to infer regulatory networks from scRNA-seq data, along with two methods for inference from bulk data (**Figure 1**, **Additional File 1**, **Table S1**). In addition to these published algorithms, we used Pearson correlation to infer a network of gene pairs with edges weighted by the experiment-wide correlation of expression. We first benchmarked these algorithms for their computing requirements, namely memory consumption and runtime, using different numbers of genes from 10 to 8000 genes (**Methods**, **Figure 2**). Of the compared methods, SCHiRM and BTR did not complete in a reasonable amount of time and were excluded from further analysis. SCRIBE and HurdleNormal could be run upto 2000 genes, however took much longer to run for more genes. Most algorithms that were runnable for upto 8k genes consumed upto 15G of RAM. Exceptions to this were SCHiRM, Inferelator and PIDC which took higher memory. SCHiRM memory consumption grew exponentially and was considered infeasible for larger gene sets. For the subsequent analysis we excluded SCHiRM, BTR and HurdleNormal.

**Figure 2.**
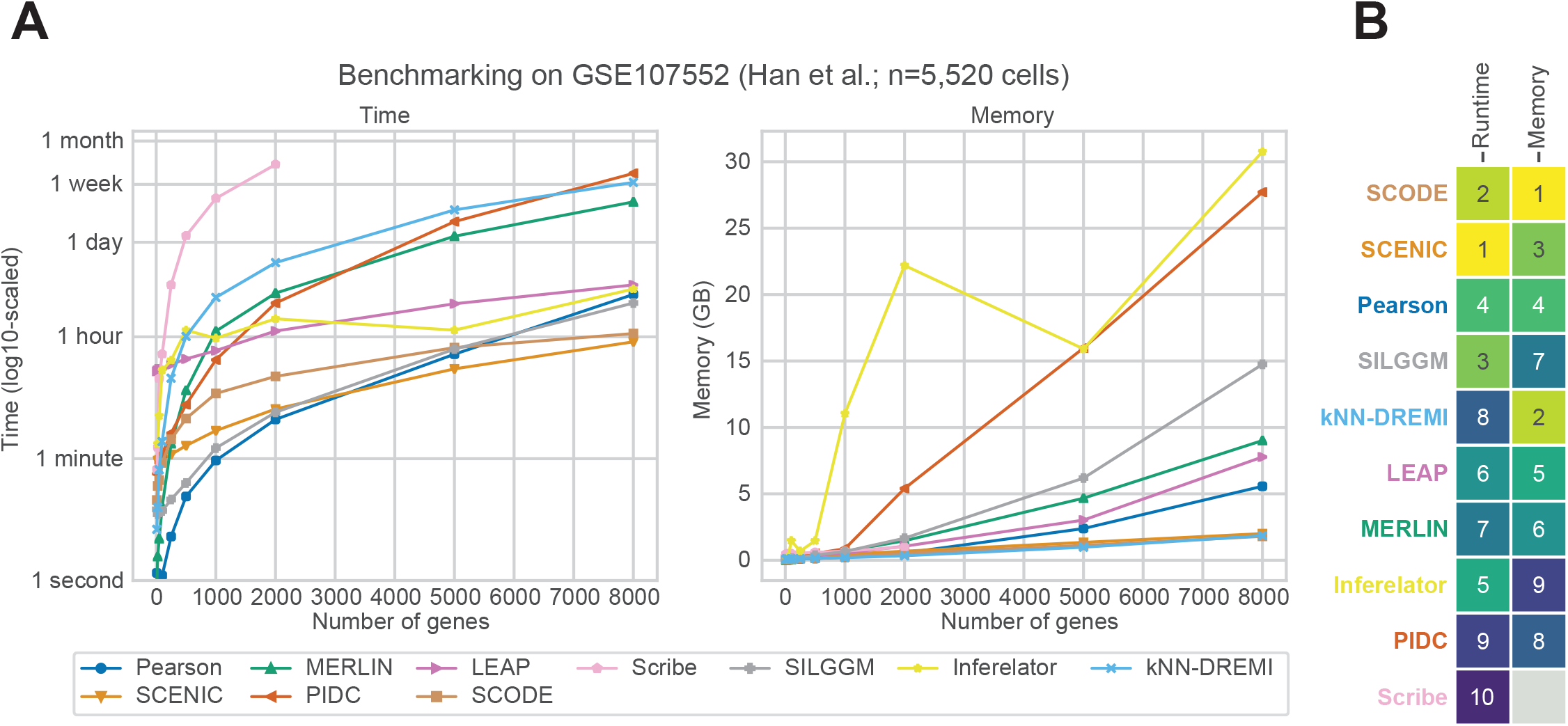
Benchmarking runtime and memory requirements. **A.** Shown are the runtime and memory requirements of each algorithm as a function of the number of genes. The x-axis shows the number of genes and the y-axis denotes the memory (left) or time (right) needed to run the algorithm. These benchmarks of resource consumption were computed using all 5,520 cells in the Han et al. dataset. **B.** Aggregated ranks of algorithms based on time and memory of algorithms. Smaller the value the better the algorithm. Some algorithms such as SCHiRM, HurdleNormal and BTR did not complete in a reasonable amount of time and were excluded from downstream analyses.

### Assessing scRNA-seq regulatory network inference algorithms based on global metrics

We next compared the performance of algorithms on gold standard networks from three species, yeast, mouse and human (**Figure 1**). These datasets included two yeast stress response datasets, four mouse datasets, one for dendritic cells, three for cellular reprogramming to pluripotent stem cells on different growth media, and one human dataset for embryonic stem cell differentiation (**Additional File 1**, **Table S2**). The datasets had a wide range of numbers of cells, ranging from 163 cells from Gasch et al [15] to the largest dataset with 36,199 cells from one of the mouse reprogramming studies [16] (**Additional File 2**). For each species dataset we had three types of gold standards, those derived from knock down or knock out of the regulator followed by global mRNA profiling (Perturb, **Figure 3**), those derived from ChIP-chip or ChIP-seq experiments (ChIP, **Figure 3**), and those derived from the intersection of these two gold standards (Perturb+ChIP, **Figure 3**). We assessed the performance of the algorithms using standard area under the precision-recall curve (AUPR) as well as F-score (**Figure 3**, **Additional File 3**, **Figure S1**) applying each algorithm on each of the datasets (**Method**). While AUPR considers all edges identified by a network inference algorithm, F-score is computed using the top edges ranked by confidence. To select the number of edges for F-score computation, we considered different number of edges and computed the stability of the metrics as a function of the number of edges (**Additional File 3**, **Figure S2**). We selected 5k edges for F-score computation as this resulted in the most stable results across algorithms.

**Figure 3.**
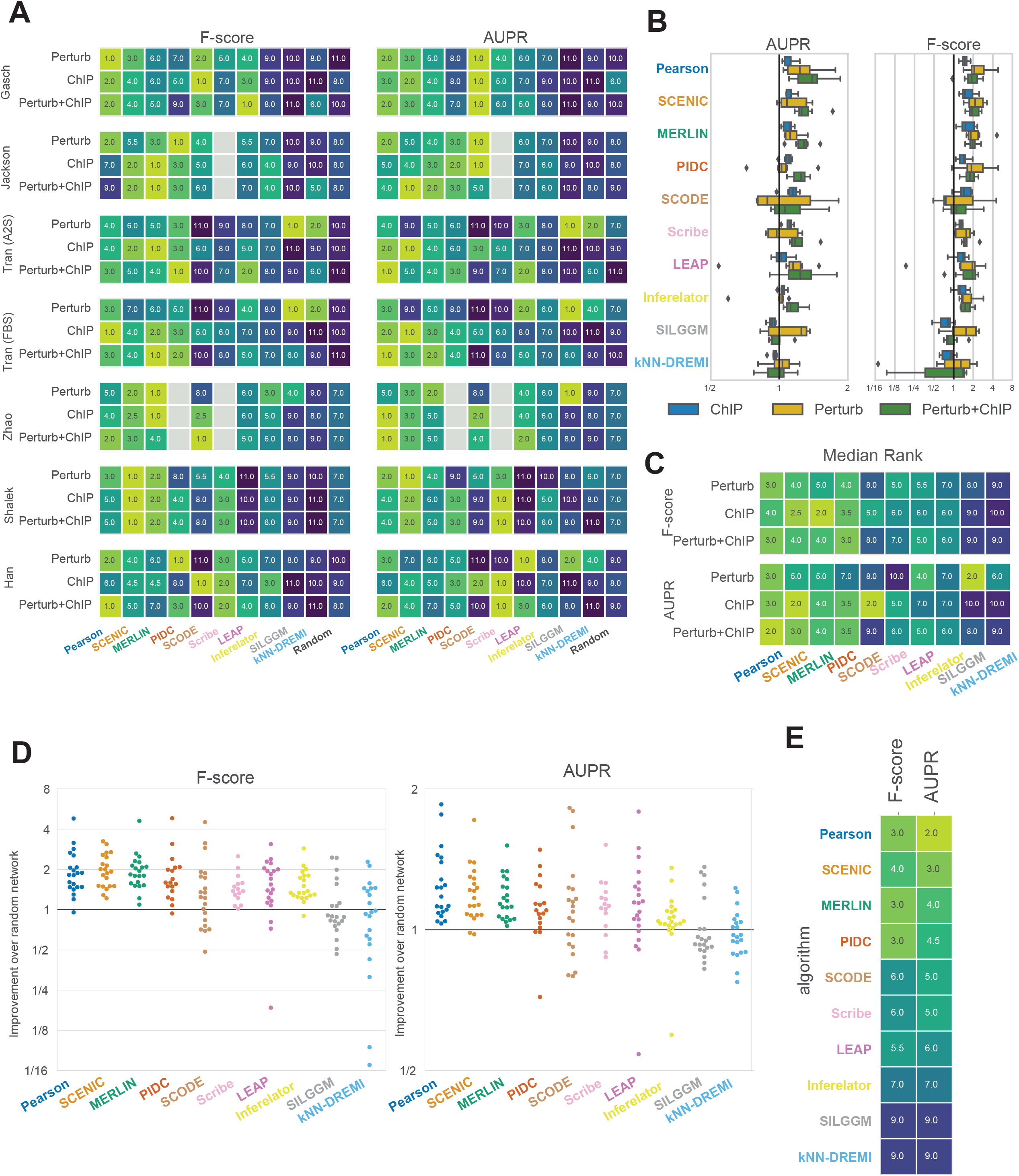
Algorithm performance across different global metrics of network structure recovery. **A.** Algorithm ranks in each of the 21 comparisons (7 datasets x 3 gold standard networks) based on AUPR and F-score. Gray cells represent the missing experiments due to scalability issue of algorithms on large datasets. **B.** Box plots showing distribution of relative performance (AUPR or F-score) compared to random (defined by a randomly weighted network with the same set of nodes and edges as the dataset) across different datasets for each type of gold standard: ChIP, Perturb and their intersection, Perturb+ChIP. The solid black line represents the random baseline. **C.** Median rank of algorithm performance with respect to each type of gold standard (median over 7 datasets). **D.** Distribution of AUPR and F-score fold improvement over random performance in the same comparison. Algorithms are ordered as shown in panel **E**. The solid black line represents the random baseline. **E.** Median rank of algorithms across all 21 comparisons using F-score and AUPR. Rows are ordered based on the mean of rank of F-score and AUPR.

We first examined the performance of each algorithm for individual datasets from the different species and ranked each method based on its F-score and AUPR (**Figure 3A**, **Additional File 3**, **Figure S1**). The ranking of methods was consistent based on both F-score and AUPR. There was more variation in ranking due to different gold standards (e.g. ChIP versus Perturb) than across datasets. Across datasets, no single method had consistently the highest scores, however the ranking of methods across datasets remained largely consistent, that is, methods that ranked high in one dataset ranked high in another dataset. A few exceptions were SILGGM and knnDREMI which had a relatively lower performance on most datasets with the exception of Tran A2S/FBS Perturb where it ranked highly. LEAP performed poorly on the Shalek dataset, but was generally among the middle ranking methods. Among the different methods, SCENIC, MERLIN and Pearson correlation were most consistently high performing across datasets based on F-score and AUPR. The size of the dataset (number of cells) was not a strong determinant of prediction performance (**Additional File 4).**

We next examined the performance of algorithms with respect to the different types of gold standards. Both F-score and AUPR were generally better for the Perturb gold standard datasets compared to ChIP gold standards (**Figure 3B**,**Additional File 3 Figure S1**). Exception was SCODE which was better for ChIP compared to Perturb. This difference in performance could be attributed to the coverage of regulator-target relationships in each datasets (**Additional File 1**, **Table S3**) and also that expression-based methods may be able to capture effects of perturbation experiments better than ChIP experiments. Difference in performance due to different gold standards was also previously observed by Pratapa et al. [13], who found poorer performance when the density of the edges was higher in a given gold standard. We aggregated the metrics across all datasets for each gold standard based on their median rank (**Figure 3C**). Based on median F-score on the Perturb gold standard, the top 3 methods were Pearson with a median rank of 3 and PIDC and SCENIC with a median rank of 4. When using ChIP, the top three methods were MERLIN, SCENIC and PIDC. Finally based on Perturb+ChIP, Pearson, PIDC were the best with a median rank of 3, followed by SCENIC and MERLIN with a median rank of 4. AUPR-based median rank varied more with the gold standard, which is reflected by overall lower correlation across gold standards ranks when using AUPR (**Additional File 3**, **Figure S3**), compared to F-score. Using Perturb, the top three methods were SILGGM, Pearson and LEAP, using ChIP the top three methods were SCENIC, SCODE and Pearson and based on Perturb+ChIP, they were Pearson, SCENIC and PIDC. Finally, we collated the metrics across gold standards and datasets (**Figure 3D**) and ranked each method based on its overall median rank across both types of gold standards (**Figure 3E**). Based on F-score, Pearson, MERLIN and PIDC had the best median rank of 3, followed by SCENIC (rank of 4) and LEAP (rank of 5.5). Based on AUPR, Pearson, SCENIC and MERLIN were the top three ranking methods.

### Assessing scRNA-seq regulatory network inference algorithms based on local network metrics

The AUPR and F-score quantify the global agreement of a predicted regulatory network with gold standard networks by comparing one edge at a time. However, from a biological application point of view, it may be beneficial to examine if there are some smaller components of the regulatory network that are predicted better than others. Therefore, we next focused on a more granular view of the inferred networks by measuring the ability to predict the targets of individual transcription factors based on the fold enrichment of true targets of a transcription factor (TF) in the predicted set as described by Siahpirani et al [17] (**Methods**). We quantified the overlap using an FDR-corrected hypergeometric test P-value and called a TF “predictable” if it had a significant P-value (FDR< 0.05). We used the number of significantly predictable TFs as another quantification of network performance (**Figure 4A**, **Additional File 3 Figure S1**). Unlike AUPR and F-score where the performance of the algorithms was often close to random, the number of predictable TFs for the random baseline was no more than 1, and typically 0. Across datasets, the methods ranked consistently, with the exception of LEAP performing much worse in Shalek and Tran (FBS) datasets, and SCODE performing much better on the Gasch and Han datasets based on the ChIP gold standard. Between Perturb and ChIP gold standards, the rankings of the algorithms were more correlated than AUPR or F-score (Pearson correlation of 0.55 for predictable TFs vs correlation of 0.41 for F-score and 0.07 for AUPR (**Additional File 3**, **Figure S3**)), but recapitulated the same preference of methods to perform better on Perturb versus ChIP datasets. We aggregated the metrics per gold standard type across datasets by computing the median rank (**Figure 4B**). Based on the Perturb gold standard, the top three methods were Pearson, PIDC and MERLIN, while based on ChIP, Peason and MERLIN were best followed by SCENIC and SCODE. Inspection of the number of predictable TFs showed there is a substantial variation across datasets and gold standards (**Figure 4C**). Finally, as in the AUPR and F-score comparison, we ranked the methods based on the predictable TFs from the three types of gold standards. Based on this ranking, the top performing methods were Pearson, SCENIC, MERLIN, and PIDC (**Figure 4D**), which was consistent with the rankings based on AUPR and F-score.

**Figure 4.**
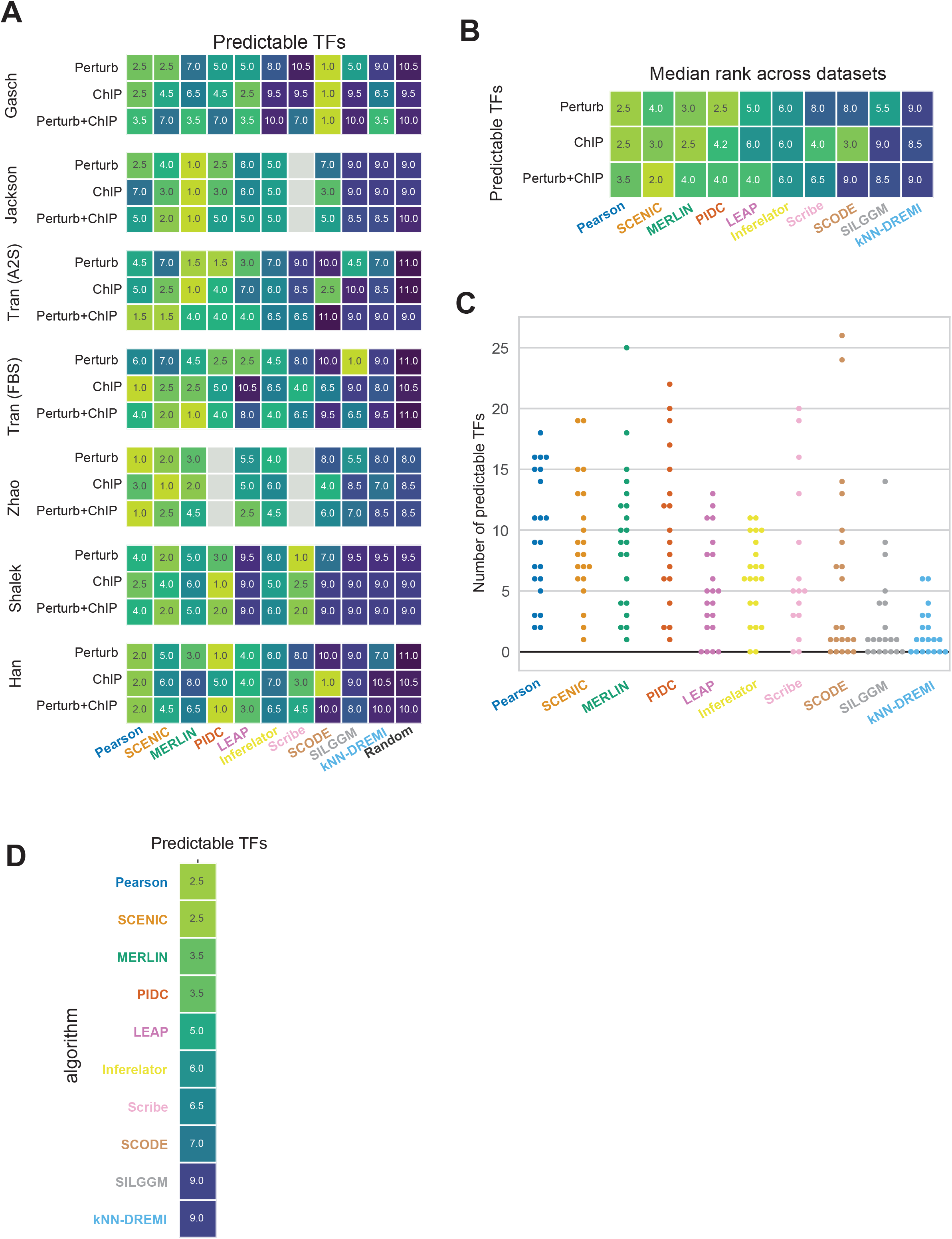
Algorithm performance based on predictable TFs. **A.** Rankings of algorithms based on the number of predictable TFs in 21 comparisons (7 datasets x 3 experimentally derived networks). Gray cells represent the missing experiments due to scalability issue of algorithms on large datasets. **B.** Median rank for each method with respect to each type of gold standard: ChIP, Perturb and Perturb+ChIP. **C.** Predictable TFs are shown as the original counts as the random network generated no predictable TFs. **D.** Overall rank of algorithms across all 21 comparisons using predictable TFs.

We next examined for each dataset which specific TFs were predicted by the different methods (**Figure 5**). This information can help determine if there are TFs that can be predicted by a specific category of methods or by all methods. For the Han dataset which profiled transcriptional programs during lineage specification from the embryonic stem cell (ESC) state, we found several ESC-specific regulators such as POU5F1, SOX2 and NANOG [18], as predictable TFs in multiple methods when using the ChIP gold-standard (**Figure 5A**). We also found TFs like CDX2 [19] and TBX3 [20], which serve as major regulators for different lineages, predicted by a number of methods. When comparing the Perturb gold standard, we found similar regulators as in the ChIP gold standard, but additionally regulators such as OTX2 and GATA3 for retinal [21] and hematopoietic lineages [22], respectively. SCODE had a number of predictable TFs in the Han dataset, however, many of them are general regulators such as SP1, YY1, POL2RA, ATF2. For the mouse cellular reprogramming datasets (Tran (FBS), Tran (A2S), Zhao), we observed a similar behavior with many of the methods identifying several developmental regulators such as Pou5f1, Esrrb and Sox2 in both ChIP and Perturb gold standards (**Figure 5B**, **Additional File 3**, **Figure S4**, **S5**). Importantly, when compared to the dendritic cell dataset (Shalek), we found a consistent over-representation of a different set of regulators across methods.This included Rel, Nfkb, Stat1 and Stat3 that are associated with immune response and were identified by several methods including SCENIC, MERLIN, PIDC, Scribe, Inferelator and Pearson (**Additional File 3**, **Figure S6**). We found a similar behavior for the yeast datasets (**Figure 5C**, **Additional File 3**, **Figure S7**), where the methods uniformly were able to recover key regulators associated with stress response such as HAP4 (oxidative stress) and GCN4 (amino acid starvation, [23]). The overall fold enrichment for predictable TFs was higher for the yeast datasets compared to the mammalian datasets. Taken together, these results show that although the performance of methods based on global metrics such as F-score and AUPR is modest, methods are able to consistently recover relevant regulators for a particular system.

**Figure 5.**
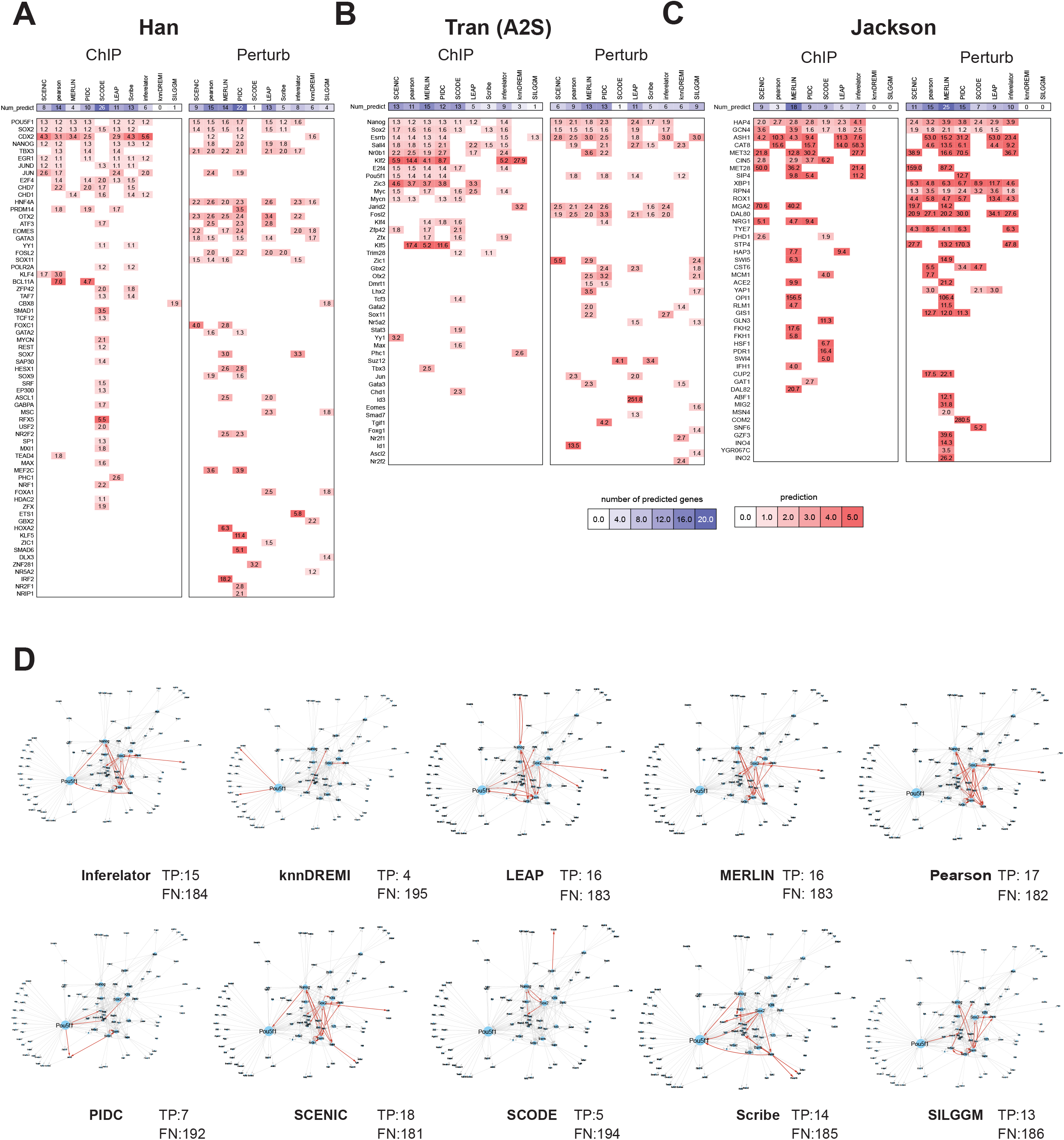
Predictable TFs identified in specific datasets. **A** Predictable TFs in the human ESC state dataset from Han et al. Rows are ordered by the number of algorithms the TF is found predictable in. The color is proportional to fold enrichment of true targets in the predicted target set of the TF. Heatmaps show the enrichment of each TF’s target set from a ChIP-based (left) or perturbation-based (right) gold standard network in an inferred network. Columns are ordered with hierarchical clustering. Grey cells indicate TFs that did not appear in one of the two experimentally derived networks. **B**. Predictable TFs in the mouse cellular reprogramming dataset from Tran et al. **C**. Predictable TFs for the yeast stress dataset from Jackson et al. **D.** Recovery of a literature curated regulatory network for the embryonic stem cell state. The network has 199 interactions curated from the literature. Shown is the overlap of this network with that inferred by each algorithm on the Tran et al. (A2S) dataset. True positive interactions are in red and false negative interactions are in grey. The number of true positives and false negatives are also listed with each method.

Finally, as large-scale gold standards themselves may be noisy, we curated a small gold standard network from regulatory interactions reported in literature for the ESC state [24–30] (**Methods**). The regulatory network has 35 regulators and 90 target genes (some of which included regulators as well) connected by 267 edges (**Additional File 2**, gold_standard_datasets.zip). We next used the top 5000 edges from the networks inferred on the Tran (A2S) dataset and depicted true positives (found by a method) and false negatives (missed by a method) among 199 of the 267 interactions that remained after removing edges due to missing expression (**Methods**). Most of the methods recovered cross regulatory interactions between Nanog, Sox2 and Pou5f1, which are master regulators for establishing the ESC state (**Figure 5D**). Methods that recovered the fewest number of true positives included SCODE, knnDREMI and PIDC. Interestingly SCRIBE which like PIDC is based on information theory was able to infer many more true positives (14) compared to PIDC (9). Overall the performance based on the recovery of true positive edges was consistent with our global and predictable TF metrics but also highlighted additional properties of the network inference methods in terms of recovery of curated regulatory interactions.

### Similarity of inferred networks from different methods depends upon the dataset

We next asked the extent to which the algorithms agree in their predictions by measuring the Jaccard similarity and F-score between the top ranked edges of each pair of inferred networks (**Figure 6**). The magnitude of the similarity increased as more edges were considered but the overall trend of pairwise similarity remained the same (**Additional File 3**, **Figure S8**, **S9**). We focused on the Jaccard scores obtained on the top 5k edges for consistency with our other results; similar behavior was observed for F-score as well (**Additional File 3**, **Figure S10**). To enable comparison of methods, we grouped the methods in two ways. First, we ordered the methods in the same way in each dataset based on the median similarity of inferred networks and visualized the pairwise similarity for each dataset (**Figure 6, Common ordering**). This allowed us to examine the change in similarity for a pair of methods as a function of datasets. Scribe’s similarity to other methods depended upon the dataset, however, it is most similar to PIDC, which is also based on information theoretic metrics. LEAP was most similar to Correlation but this also depended upon the dataset; e.g. high Jaccard score in Gasch, Jackson and Zhao and low scores in other datasets. Some datasets resulted in more similar networks compared to others. In particular, methods were most similar (high Jaccard coefficient) in the Han and Jackson datasets and were least similar for the Shalek and Gasch. These two datasets were also the smallest of the two datasets, which could explain the lower similarity. Methods that tended to have the highest similarity with other methods were Pearson, SCENIC, Inferelator and PIDC.

**Figure 6.**
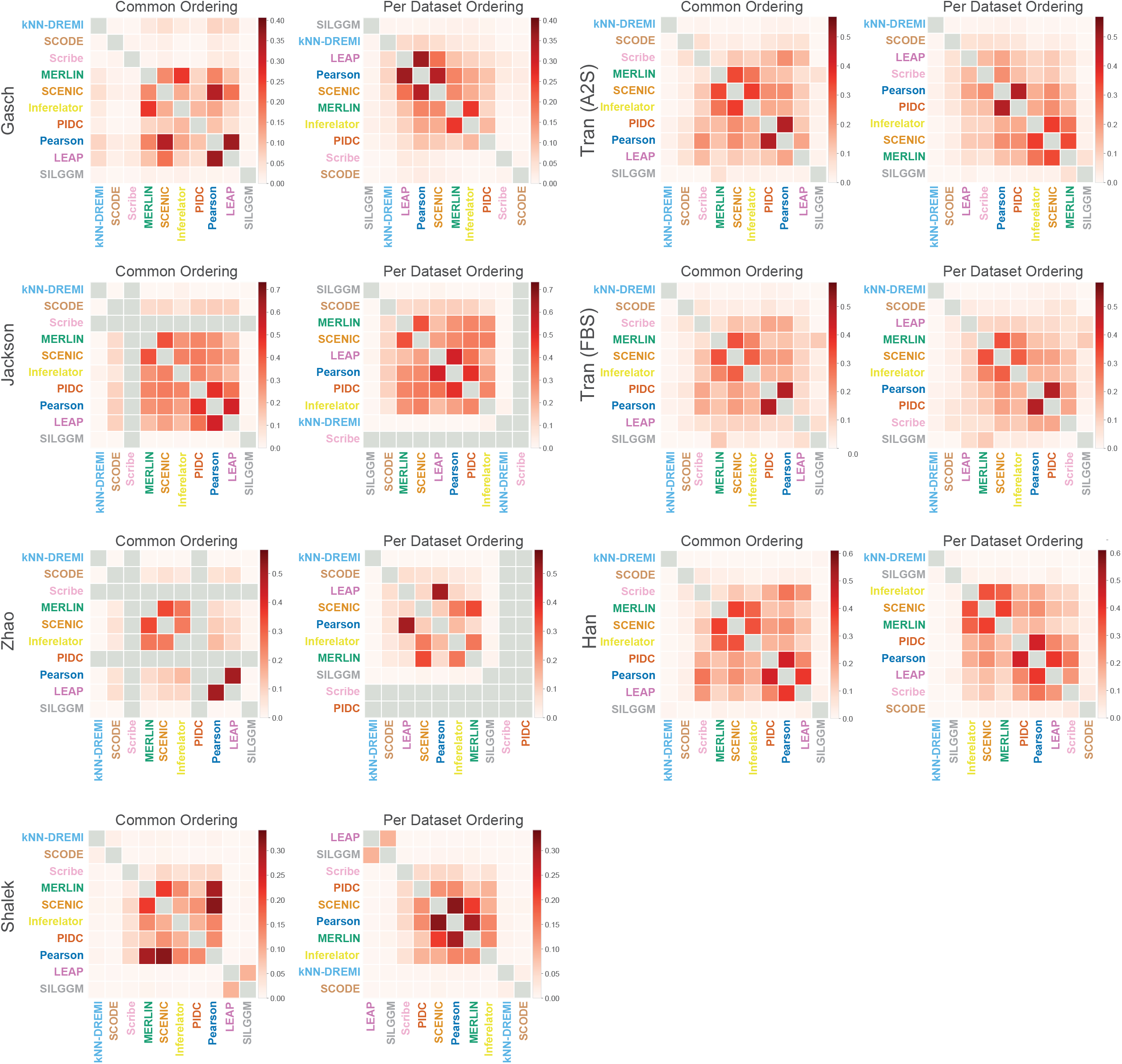
Pairwise network similarity between inferred networks for each method. Shown here are the Jaccard similarity between the top 5,000 edges in each pair of networks inferred on each dataset. **Common Clustering.** The columns and rows of each matrix are ordered with respect to a hierarchical clustering of the median similarity across all datasets. **Per Dataset clusters** The columns and rows are ordered with respect to each dataset.

We next ordered the methods based on the Jaccard similarity per dataset producing dataset-specific orderings (**Figure 6, Per Dataset Ordering**). SILGGM and knnDREMI generally learned different networks compared to other methods, including those based on the same class of models, e.g. information theory for knnDREMI (PIDC, Scribe) and probabilistic models for SILGGM (MERLIN, Inferelator). The methods that exhibited the highest similarity across datasets were LEAP, PIDC and Pearson, with the exception of Tran (FBS), Tran (A2S) and Shalek where LEAP learned more different networks and Gasch where PIDC was different from Pearson and LEAP. This was followed by SCENIC, MERLIN and Inferelator, which also tended to group together in multiple datasets. In summary, these comparisons revealed that the similarities between methods depended upon the dataset, however, several of the methods such as PIDC and Correlation tended to infer similar networks across datasets. Furthermore, the networks inferred from the same family of algorithms were not necessarily more similar than from different families of algorithms.

### Imputation of scRNA-seq datasets does not improve network inference

Single-cell expression data is characterized by a large number of zero expression counts which can arise due to technical (low depth) or biological reasons (cell type-specific expression). To address this problem, several methods have been developed to impute the value of missing expression counts [31]. We applied the MAGIC algorithm to each dataset [4], which was one of the top imputation methods from a recent benchmarking study [31]. MAGIC computes pairwise similarity between cells and creates a Markov transition graph based on these similarities. It then “diffuses” expression counts among similar cells based on this Markov transition graph. We inferred networks using the imputed data from each experiment and compared their AUPR, F-score and predictable TF metrics with those of the networks inferred from the original sparse datasets (**Figure 7**). Based on F-score (**Figure 7A**), most methods did not benefit from imputation across datasets. An exception to this were the Shalek and Gasch datasets, which as noted before are among the smaller datasets. We examined the ratio of F-score after and before imputation across different gold standards and datasets and found SCODE and kNN-DREMI to improve when using imputed data (**Figure 7B**). However, imputation generally did not benefit the network inference procedure with most algorithms exhibiting a ratio < 1 across datasets and gold standards. We repeated these comparisons based on AUPR (**Figure 7C, D**) and predictable TFs (**Figure 7E, F**). AUPR did not change much with imputation, however, we noted a similar boost in performance for the Shalek dataset. Finally, based on predictable TFs, performance generally did not improve with the exception of the Tran (A2S), Tran (FBS) and Shalek datasets where algorithms such as SCODE and knnDREMI showed a benefit (**Figure 7E, F**). Taken together, our experiments across different datasets and gold standards showed that imputation benefitted in a handful of cases of algorithms and datasets, but in the majority of the cases either stayed the same or had a detrimental effect.

**Figure 7.**
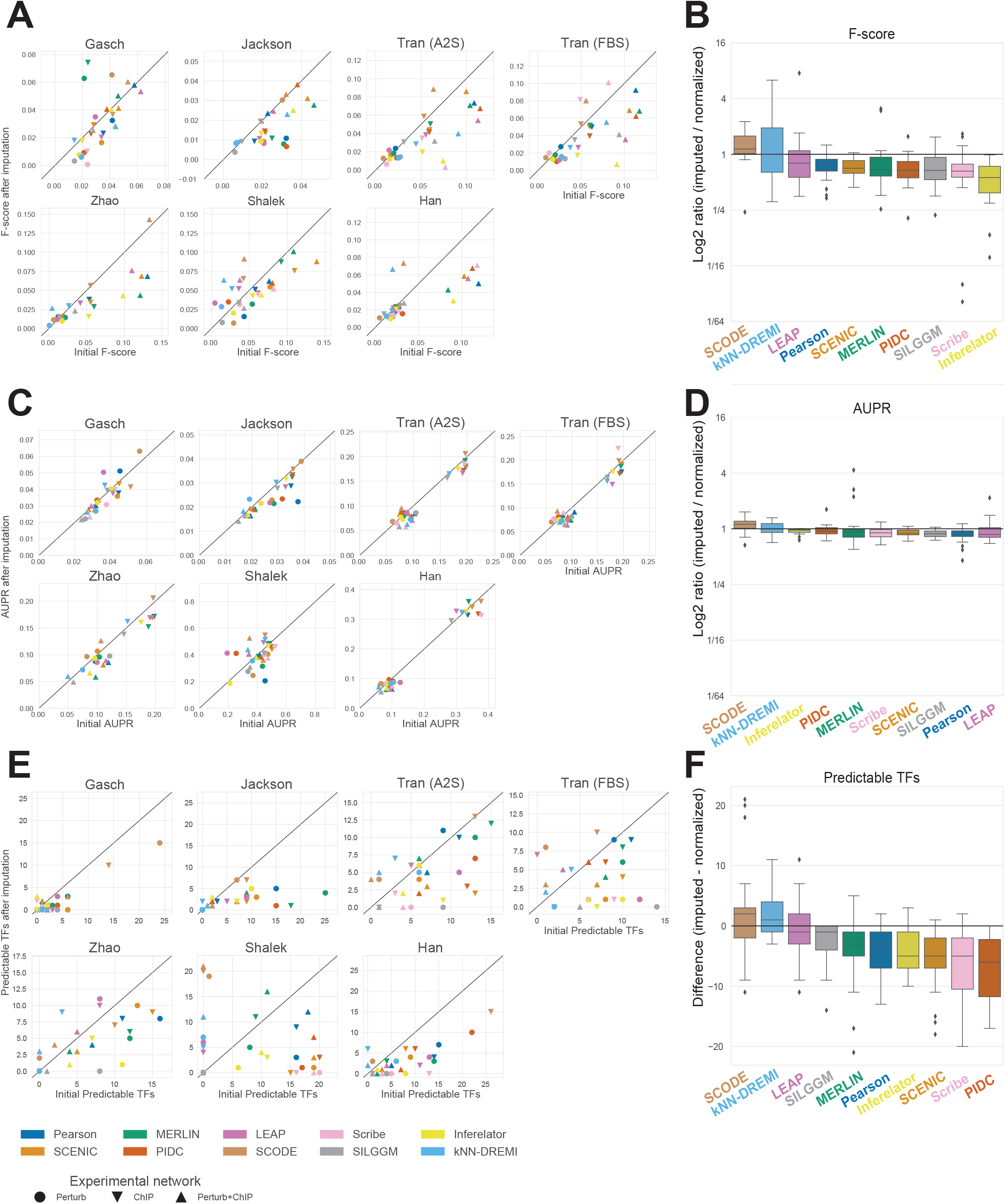
Effect of imputation on inferred networks. **A.** Each separate plot corresponds to a dataset, the shape of the marker denotes the type of gold standard and the color depicts the method of network inference. F-score before imputation is on the x-axis and F-score after imputation is on the y-axis. A marker above the diagonal corresponds to improved performance after imputation. **B.** The box plots summarize the relative performance after imputation compared to before imputation over the 21 comparisons: 7 datasets x 3 gold standard networks. Algorithms are ordered based on median performance gain. **C.** Similar to **A**, showing AUPR. **D.** Similar to **B** reporting relative AUPR after and before imputation. **E.** Similar to **A**, showing the number of predictable TFs. **F.** Similar to **B**, reporting the difference in the predictable TFs between the two networks after and before imputation.

### Comparison of network inference from single cell versus bulk expression datasets

Before the availability of scRNA-seq datasets, expression-based network inference was performed with bulk expression datasets, which required large sample sizes. Such datasets are available from a single study in yeast [23] or can be created by collecting studies from public gene expression omnibus. Among the conditions (species and cell states) we examined in this study, we had matching bulk expression datasets in yeast (stress conditions), mouse ESC (mESC) and human ESC (hESC). We therefore applied network inference on these bulk expression datasets using five state of the art algorithms and compared the performance of the methods based on the same gold standards and metrics of F-score, AUPR and predictable TFs (**Figure 8**, **Methods**). For the scRNA-seq comparison, we used the inferred networks from the Gasch scRNA-seq dataset for yeast, Tran (A2S) for mESC and Han dataset for hESCs.

**Figure 8.**
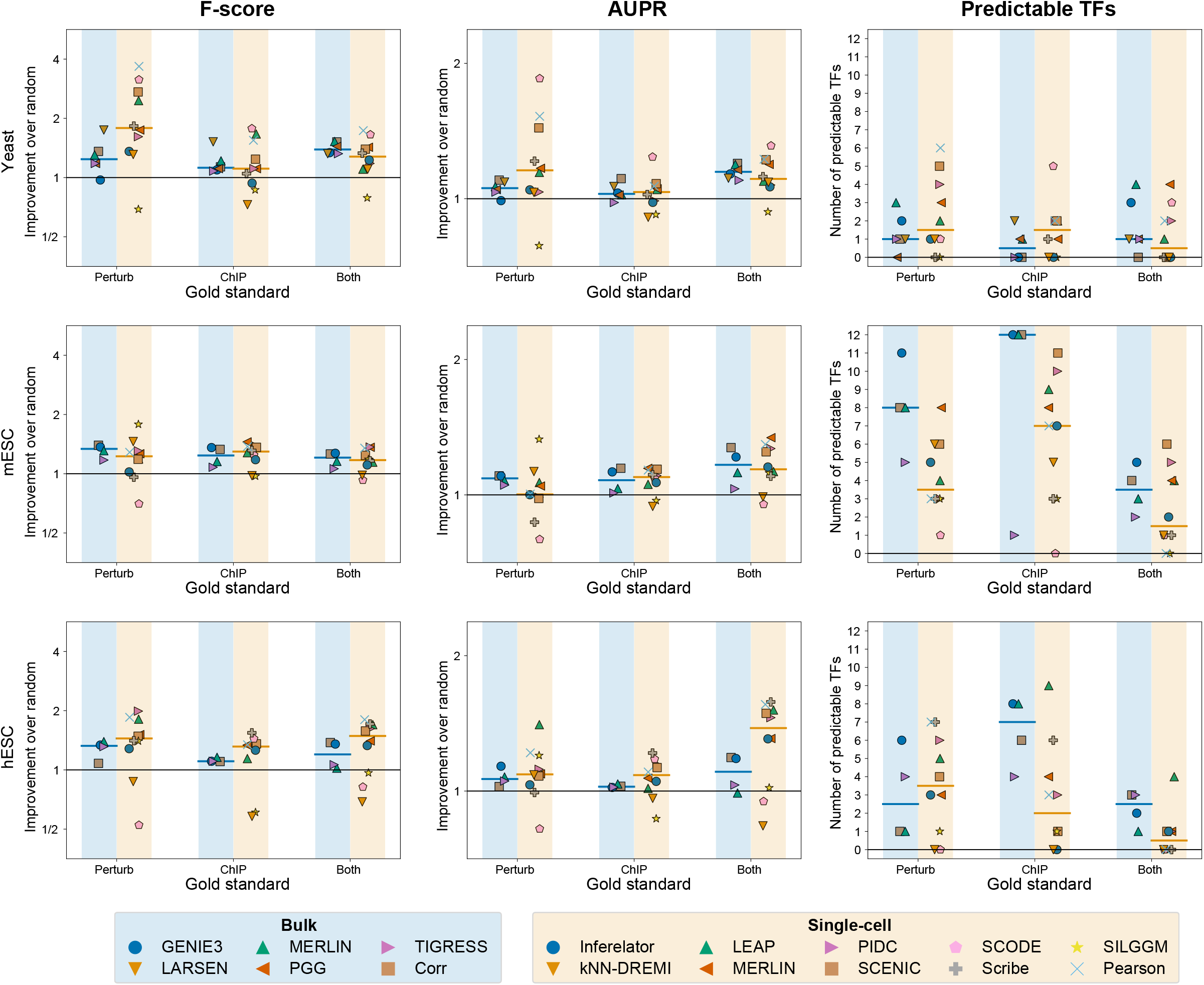
Comparison of network inference on bulk versus single cell RNA-seq data. Shown are the relative F-score and AUPR scores compared to random and the number of predictable TFs on comparable bulk and scRNA-seq datasets. Methods used for network inference for bulk and single cell datasets are listed at the bottom. Each marker on each plot corresponds to a method for bulk (blue shading) or scRNA-seq data (yellow shading). The yeast dataset compared the scRNA-seq data from Gasch et al. [15], to the bulk dataset from the same author [23]. Human and mouse bulk datasets were generated from publicly available datasets (**Additional File 1**, **Table S3**).

Based on F-score scRNA-seq-based networks exhibited a moderate improvement on the Han (human ESC) dataset and yeast dataset when using either the Perturb+ChIP or Perturb gold standards respectively. The results were largely consistent for AUPR and F-score. However, when comparing predictable TFs, we observed a marked decrease in the number of predictable TFs when comparing the results for mouse ESC and human ESC datasets using ChIP and ChIP+Perturb. Overall this suggests that individual scRNA-seq datasets capture meaningful variation that offers comparable performance to bulk expression datasets collected from a large number of experiments.

### Addition of priors and transcription factor activities can improve performance

Our experiments so far utilized gene expression alone for network inference. Bulk network inference methods that incorporate additional genomic sources such as sequence-specific motifs of TFs have improved performance [17,32]. Such motif-based TF target relationships have been used as priors to constrain the graph structure [17,32] and also estimate TFA [33–35], which have both led to improvements in network recovery. We therefore examined if incorporation of prior knowledge is beneficial for network inference from scRNA-seq datasets. We created graphs encoding prior knowledge of regulator-target relationships as described in [17] (**Methods**). We considered MERLIN, Inferelator and SCENIC, because only MERLIN and Inferelator incorporate priors, and SCENIC was among the top performing methods with efficient run times. We estimated the TFA of the regulators based on the structure of the prior network using network component analysis (NCA) method [33]. Subsequently, the estimated TFA of these regulators along with their gene-level expressions were utilized to find their target genes. For Inferelator we estimated the performance using NCA-based TFA and its own TFA estimation, while MERLIN and SCENIC used the NCA-based TFA (Referred to as NTFA). We add “+P” after an algorithm’s name if it used prior information and “+TFA” or “+NTFA” if it used TFA. We applied these algorithmic configurations to the Gasch, Jackson, A2S, FBS, Shalek and Han datasets and assessed performance using the ChIP, Perturb and Perturb+ChIP gold standards. We did not include the Zhao dataset due to the computational burden of this dataset. MERLIN and Inferelator were evaluated for the ability to incorporate prior knowledge and TFA, while SCENIC was evaluated for its ability to leverage TFA. We compared these algorithms to their versions that did not incorporate TFA or prior and to the baseline Correlation (**Figure 9**, **Additional File 3 Figure S11**).

**Figure 9.**
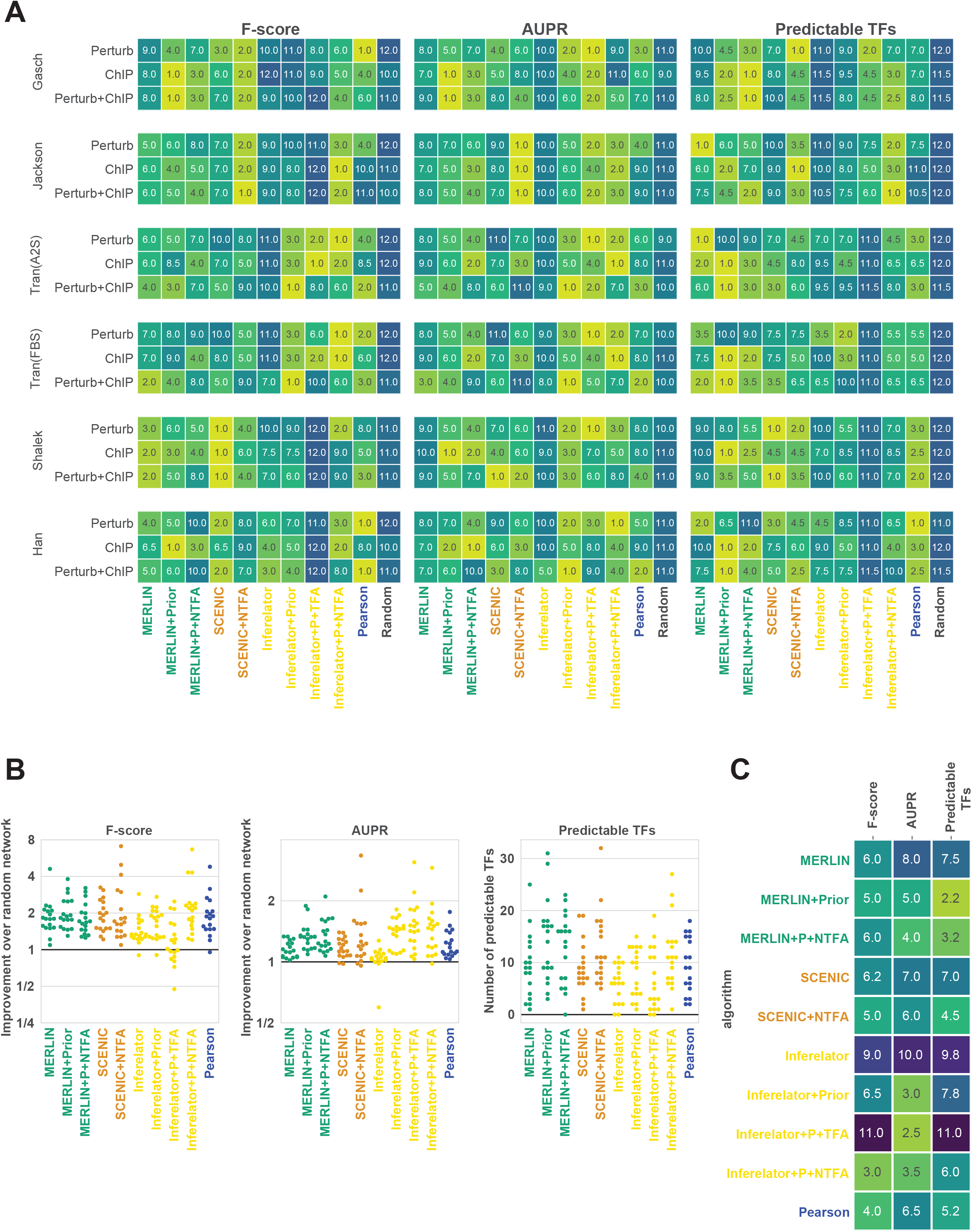
Assessing the utility of incorporating prior knowledge and TFA in network inference on scRNAseq data. **A.** Ranks of each method based on F-score, AUPR and predictable TFs shown for 6 datasets using the three gold standards. Methods compared include MERLIN, Inferelator and SCENIC and their enhanced versions for prior or TFA. The terms “+P” after an algorithm’s name denotes adding prior while addition of “+TFA” or “+NTFA” corresponds to TFA estimated using Inferelator’s inbuilt TFA estimation or TFA estimated from Network Components Analysis (NTFA), respectively. MERLIN+NTFA, MERLIN+Prior, MERLIN+P+NTFA, are extensions of MERLIN using NTFA, Prior, and Prior and NTFA, respectively. SCENIC+NTFA, uses NTFA-based TF activity in addition to gene expression. Inferelator+Prior, Inferelator+Prior+NTFA, Inferelator+Prior+TFA were extensions to Inferelator for Prior or Prior and TFA. Also included for comparison are the network inference results for Pearson correlation and a random network. **B.** Distribution of fold enrichment of F-score or AUPR over random or the number of TFs across the datasets. **C.** Aggregated ranks of the methods across datasets and gold standards.

Based on F-score, MERLIN with prior (MERLIN+P) outperformed MERLIN (8 out of 18 times, **Figure 9A**, **Figure S11**), and incorporation of TFA (MERLIN+P+NTFA) outperformed MERLIN alone for 8 out of 18 of the dataset and gold standard combinations. Although MERLIN+P and MERLIN+P+NTFA outperformed MERLIN in less than 50% of the comparisons, they had better fold enrichments (**Figure 9B**) and overall better rank (**Figure 9C**). SCENIC with TFA (SCENIC+NTFA) outperformed SCENIC alone for 9 out of 18 dataset and gold standard combinations. The biggest gain for using priors was observed with Inferelator with Inferelator+P outperforming Inferelator for 11 out of 18 of the dataset and gold standard combinations. Incorporation of TFA in Inferelator+P+TFA led to higher performance in 7 out of 18 cases. Finally addition of NCA based TFA and prior was most beneficial making Inferelator alone to be outperformed in 15 out of 18 of combinations. Finally, when examining F-score ranks of Pearson, methods that used priors had a higher overall rank than Pearson correlation (**Figure 9B, C**). Based on AUPR, we found a greater benefit of methods using priors or TFA. In particular, MERLIN+P and MERLIN+P+NTFA outperformed MERLIN in 17 and 15 out of 18 dataset and gold standard comparisons, respectively. SCENIC+NTFA outperformed SCENIC in 11 out of 18 comparisons. Inferelator+P, Inferelator+P+TFA, Inferelator+P+NTFA consistently outperformed Inferelator in the majority of the comparisons. Compared to Pearson correlation nearly all methods that used prior or TFA outranked Pearson correlation (**Figure 9C**). Finally, based on the number of predictable TFs, MERLIN+P and MERLIN+P+TFA outperformed MERLIN in 12 and 13 out of 18 cases, respectively. SCENIC+NTFA outperformed SCENIC in 11 out of 18 cases, and Inferelator+P, Inferelator+P+TFA and Inferelator+P+NTFA outperformed Inferelator in 12, 6, 11 out of 18 cases respectively. As in the case of AUPR and F-score Pearson correlation was outperformed in overall rank by a method that used priors (MERLIN+P, MERLIN+P+NTFA) or TFA (SCENIC+NTFA, MERLIN+P+NTFA). Taken together, these results show that inclusion of prior information to constrain graph structure or for TFA improves the quality of inferred networks for scRNA-seq datasets as well.

## Discussion

The rapid growth of scRNA-seq datasets has opened up remarkable opportunities in the field of expression-based gene regulatory network inference. Accordingly, substantial effort has been invested to develop and apply regulatory network inference algorithms to single cell datasets. Here we benchmarked the computing requirements and overall performance of methods in network structure recovery across a large number of datasets spanning different species and model cell lines. Our work expands on previous efforts by considering larger scRNA-seq datasets across multiple model systems, additional classes of methods, different types of metrics and gold standards that can provide biological insight into the predictions made by the different methods.

We evaluated methods based on their computing requirements as well as performance on different gold standard networks. Due to excessive computing requirements we could not rank several methods that were specifically geared towards handling the statistical properties of single cell datasets, such as SCHiRM, HurdleNormal and BTR. As single cell datasets grow, efficient implementation of algorithms would be important. Our evaluation metrics included both global metrics such as AUPR and F-score as well as local metrics such as the number regulators whose targets could be predicted significantly. Based on AUPR and F-score, the overall performance of methods remains modestly better than random, however, we find predictable TFs as a more sensitive metric that highlights the strengths of network inference methods. Importantly, different methods were able to recapitulate relevant regulators for the system of interest, for example key stem cell regulators in the developmental datasets and immune response regulators in the dendritic cell dataset. No single method was a winner across all datasets and gold standards, however, based on the overall performance and computing requirements of the methods, the methods PIDC, SCENIC, MERLIN and Pearson are the top methods. Beyond using expression alone, we assessed the benefit of using prior networks for both constraining the network structure [17,32] as well as estimating hidden TFA [35]. We found that incorporation of sequence motifs as priors to constrain the network structure or to incorporate TFA during network inference is beneficial for improved recovery of networks from scRNA-seq data. Importantly, methods that used priors or TFA had better overall performance. Our results suggest that future efforts for transcriptional network inference from scRNA-seq datasets must leverage sequence motifs. The availability of relevant single cell or bulk accessibility datasets could greatly benefit the generation of high quality network priors and ultimately the gene regulatory networks integrating these priors with expression.

One challenge with scRNA-seq datasets is the high proportion of zeros, which can be due to both biological as well as technical reasons. As imputation has been proposed as an approach to handle sparse datasets such as scRNA-seq datasets, we studied the impact of imputation on the quality of our inferred networks. As such imputation did not improve the performance of most methods, however, the datasets where it did benefit tended to have relatively smaller number of cells. One caveat in our analysis is that we considered only one imputation method, MAGIC, which was shown to be one of the top imputing methods. A direction of future work is to consider additional imputation methods to examine the impact of imputation on network inference. Related to the depth issue, we briefly explored the effects of differences in cell counts of datasets on network inference. We found that the performance of a method was not necessarily similar on two distinct datasets simply because the datasets have the same number of cells. In particular, we observed that decreasing the number of cells in Jackson (17,396 cells) to match the Gasch scRNA-seq dataset (163 cells) or Zhao (36,199 cells) to Tran (FBS) size (3,324 cells) did not have the same effect. The performance stayed largely unchanged for Zhao, but decreased for Jackson. One of the factors could be the difference in the sequencing depths of the datasets. We observed that the total number of transcripts (across all genes) per cell and the number of distinct genes detected per cell are higher in Gasch compared to that of the Jackson subsamples (see Supplementary Text). It supports the observation that the methods utilized in the study (SCENIC and MERLIN) consistently achieved higher AUPR and F-scores with Gasch compared to that with the Jackson subsamples. On the other hand, for Tran (FBS) vs. the Zhao subsamples, the depth is more similar and the change in the number of cells did not affect the performance (see **Supplementary Text 1**). These results suggest that depth rather than number of cells in the dataset are important considerations when comparing results across datasets. Our observations also showed that SCENIC was less susceptible to dataset size change than MERLIN, which would indicate that the non-linear, non-parametric model of the Random Forests regression is favorable for modeling scRNA-seq profiles. To handle low depth data might be use pseudobulk expression of a small cluster of cells, e.g. “metacells” [36], which were shown to be advantageous for a number of analytical tasks for scRNA-seq datasets [37]. A direction of future work would be to examine the effect of metacells on computational costs and accuracy of network inference algorithms.

Single cell transcriptomic datasets have the advantage that a single experiment can produce large number of samples that are comparable or larger than existing bulk datasets that have been used for network inference. Therefore, we compared the quality of the inferred networks from bulk and scRNA-seq datasets for yeast, mouse and human using methods for both bulk and single cell datasets. We find that scRNA-seq datasets, despite being sparse are able to capture sufficient meaningful biological variation and perform at par when using bulk RNA-seq datasets. Our study however is not perfect since the bulk and single cell datasets were collected from different sources. Generating controlled datasets capturing bulk and single cell profiles for the same system could provide additional insight into the relative advantage of scRNA-seq datasets for inferring gene regulatory networks.

The gold standards and datasets that we have collected in our work should be beneficial for these future studies. Another direction of future work is to leverage the inherent heterogeneity and population structure of scRNA-seq datasets. Methods based multi-task learning is a promising framework to model population and network heterogeneity [38,39]. The relatively good performance of a simple Pearson correlation when using expression alone was surprising. This could be an artifact of our gold standards which are admittedly imperfect. Correlation could remain as a useful baseline for network inference method development, however, it does not permit the incorporation of prior knowledge which we found to be crucial for obtaining the best performance in network inference. Generation of improved gold standards, especially for mammalian systems, based on newer high throughput perturbation studies such as Perturb-seq [40] could significantly benefit our ability to infer genome-scale gene regulatory networks.

## Methods

### Dataset preprocessing

We obtained a total of seven scRNA-seq datasets from three different species, human, mouse and yeast (see **Additional File** 1, **Table S1** for details). Each dataset was filtered in a two step process. First, we filtered out genes that were expressed (count >0) in fewer than 50 cells and cells with fewer than 2000 total UMIs. Next, we added back regulatory proteins such as TFs and signaling proteins for each dataset in which they were expressed. Each dataset was depth normalized and the count was square-root transformed. Finally we removed genes that were not measured in any of the gold standards associated with the dataset. Preprocessing scripts are available at https://github.com/Roy-lab/scRNAseq_NetInference/tree/master/scripts/data_preprocessing. For performing the memory and time profiling experiments, we used the Han human pluripotent stem cell dataset [41], which had a total of 5,520 cells and selected number of genes, *n* ∈ {10, 25, 50,100, 250, 500,1000, 2000, 5000, 8000}. Each algorithm was applied on each dataset and profiled for memory requirements and execution time. Runtime and memory were captured using the `/usr/bin/time -v` command with Linux.

### Regulatory network inference algorithms for single cell RNA-seq datasets

To assess algorithms for regulatory network inference, we considered recently published methods that span a variety of modeling formalisms including ordinary differential equations, probabilistic models and information theoretic methods. We considered algorithms developed for both bulk and scRNA-seq datasets for network inference. Here we provide a brief summary of each algorithm included in our analyses (see **Additional File** 1, **Table S2** for more details about the algorithm implementation).

#### BTR

BTR is based on a Boolean model representation of a gene regulatory network [7]. Here a regulatory interaction is inferred by assuming a gene has a value of either 0 or 1 and the state of a target gene is specified by a boolean function of the state of its regulators.

#### HurdleNormal

This approach is based on a multi-variate Hurdle model which is a probabilistic graphical model [12] suited for handling zero-inflated datasets. The graph structure, which represents the gene regulatory network is estimated by first defining the model in terms of a set of conditional distributions and solving these regression problems by solving a set of penalized neighborhood selection problems.

#### scHiRM

Single cell hierarchical regression model (SCHiRM) is a probabilistic model based framework suited for handing sparse datasets such as those from scRNA-seq experiments [11]. The expression level of a gene is modeled as a Poisson log distribution with overdispersion. A regulatory network is estimated by a regression function that links the expected expression level of a gene to the expected expression level of target genes using a hierarchical framework.

#### kNN-DREMI

kNN-DREMI recovers gene-gene relationships by adapting the DREMI algorithm [42] to scRNA-seq data [4]. The DREMI makes use of a conditional density to estimate mutual information between pairs of genes. It makes use of a heat-diffusion based kernel density estimator to estimate the joint density, followed by conditional density estimation. kNN-DREMI replaces the heat diffusion kernel with a knn-graph based density estimator to handle the high-dimensionality and sparsity of the scRNA-seq data.

#### LEAP

LEAP [10] requires cells be ordered along a pseudotime trajectory, and computes pairwise correlation between genes *i* and *j* at various lags along this trajectory. The algorithm chooses a window size *s*, and for all possible start times *t* and lags *l*, computes the Pearson’s correlation coefficient between the first s observations of *i* beginning at time *t* and the s observations of *j* beginning at time t + l. The score for a regulatory relationship from *i* to *j* is the maximum of all computed correlation coefficients *pjj*.

#### PIDC

PIDC [2] is based on an information-theoretic framework and estimates a specific type of multivariate information called Partial Information Decomposition (PID) between three random variables, each representing a gene. PID is defined using the “unique”, “synergistic” and “redundant” information between two variables, *X* and *Y* and a third variable *Z*. The “unique” information is the contribution to the multiinformation specifically from two of the variables, “redundant” is the contribution from either *X* or *Y*, while “synergistic” is the contribution from both *X* and *Y* to *Z*. In the context of network inference, PIDC makes use of the ratio of the unique information to the mutual information between two variables and establishes an edge based on the magnitude of this ratio.

#### SCENIC

SCENIC is based on the GENIE3 [43] network inference algorithm, which infers a gene regulatory network by solving a set of regression problems using ensemble regression trees. SCENIC also offers two additional steps, RcisTarget for enriched motif identification in the target set of a TF and AUCell to determine the activity of a set of targets of a TF in each cell. For our experiments we only used the GENIE3 outputs from SCENIC.

#### SCODE

SCODE is based on an Ordinary Differential Equation based formulation of a gene regulatory network and requires pseudotime as input in addition to the expression matrix for learning the regulatory network [6]. SCODE makes use of a lower dimensional representation of the major expression dynamics of regulators, **z** and uses a linear transformation matrix, **W** to estimate the observed expression dynamics of all genes from **z**. The gene regulatory network is represented as a function of **W** and estimated by solving a set of linear regressions.

#### Scribe

Scribe [3] is based on an information theoretic framework that uses pseudotime as an additional input for regulatory network inference. Scribe is based on a specific type of information measure called Restricted Directed Information that measures the mutual information between the current state of a target gene and a previous state of a regulator, conditioned on the previous state of the target gene. Scribe additionally uses a bias correction scheme of the samples such that cells capturing transitioning states are weighted more towards establishing the regulatory relationships.

#### SILGGM

SILGGM makes use of a Gaussian graphical model [1] representation of a gene regulatory network and offers several recent and efficient implementations for estimating conditional dependence between random variables (genes): bivariate nodewise scaled Lasso, de-sparsified nodewise scaled Lasso, de-sparsified graphical Lasso, and GGM estimation with false discovery rate (FDR) control using Lasso or scaled Lasso. For our experiments, we applied SILGGM with de-sparsified nodewise scaled Lasso implementation of neighborhood selection, which is the default algorithm for SILGGM.

#### Bulk dataset network inference methods

In addition to above methods which are specifically geared for scRNA-seq datasets, we also considered methods developed for bulk RNA-seq datasets: MERLIN [44] and Inferelator [32]. MERLIN is based on a dependency network learning framework and employs a probabilistic graphical model prior to impose modularity prior. We ran MERLIN with stability selection and default parameter settings for the sparsity *p* = –5, modularity *r* = 4, and clustering *h* = 0.6. Inferelator is also based on a dependency network learning framework which uses regularized regression for network inference. We used the BBSR algorithm within Inferelator, v0.3.0 downloaded from https://github.com/simonsfoundation/inferelator_ng and used Inferelator’s internal stability selection to obtain edge confidence.

#### Pearson and random networks

Our analyses included two inferred networks as baselines. The “Pearson” network was an undirected fully connected network, with edges weighted by the Pearson correlation between each pair of genes over all cells. The “random” network was an undirected fully connected network, with edge weights drawn from a uniformly distribution Unif(0,1).

### Comparison to network inference on bulk expression data

Methods used to infer networks on bulk data include GENIE3 [43], TIGRESS [45], MERLIN [44]. To run GENIE3 we used *K* = `sqrt` and `nb_trees` = 1000, to run TIGRESS we used R=1,000, α=0.3, and L=30, and to run MERLIN we used default configuration of sparsity *p =* –5, modularity *r* = 4, and clustering cut-off *h =* 0.6. MERLIN was run in a stability selection framework, where 100 subsets were created by randomly selecting half the samples in the input expression matrix, a network was inferred on each subset, and a consensus network was made by calculating the frequency of observing each edge in the 100 inferred networks. As in the single cell case, we created a network where edges were ranked by absolute value of Pearson correlation of expression profile of genes. PGG (**P**er **G**ene **G**reedy) is MERLIN without modularity prior and was run using the same parameters and in a stability selection framework similar to MERLIN. To run LARS-EN we used `lambda2`= 1E-6 and the L1 penalty was set in a 10 fold cross validation. We ran LARS-EN in a stability selection procedure similar to MERLIN. We used the MATLAB implementation of LARS-EN from the Imm3897 package downloaded from http://www2.imm.dtu.dk/pubdb/views/publication_details.php?id=3897.

We performed these comparisons on three datasets that came from comparable conditions for bulk and single cell. In yeast, we used the environmental stress response data from Gasch et al [23] for bulk to match our scRNA-seq dataset (Gasch) also generated by Gasch and colleagues [15]. For human and mouse, we curated publicly available datasets from the gene expression omnibus (**Additional File 1**, **Table S4**) for human and mouse embryonic stem cell state (Siahpirani in preparation). We compared the performance of bulk human dataset to that from the Han dataset [41] and the performance of the bulk mouse dataset to that from the Tran (A2S) dataset [46]. Because the gene sets of the bulk and single cell data were different, we filtered the gold standards to include TFs and targets that were in both datasets to make the results comparable.

### Assessment of the inclusion of priors and TFA on network inference

We tested the addition of priors and TFA on network inference performance for the MERLIN, SCENIC, and Inferelator algorithms. Specifically, for both MERLIN and Inferelator we tested the addition of priors, and priors with TFA. For Inferelator we used the TFA estimated using the NCA algorithm [33] (referred as NTFA) as well as its inbuilt TFA estimation. For SCENIC we tested the addition of TFA. We selected these algorithms because they were either among the top performing algorithms and/or the ability to incorporate this additional information. We referred to these extended versions of each algorithm with “+P” for addition of prior (e.g. MERLIN+P was MERLIN with prior knowledge), “+TFA” or “+NTFA” for methods using TFA (e.g. SCENIC+NTFA) and “+P+TFA” if the algorithm used both prior and TFA (e.g. Inferelator+P+TFA). MERLIN+P and MERLIN+P+NTFA was applied in a stability selection framework using the same 100 subsets utilized for the algorithm without the use of additional information. SCENIC and Inferelator perform internal bootstrapping to extract confidence estimates on edges. We applied the algorithms to all of the datasets utilized in this paper with the exception of Zhao due to the size of the dataset and time constraints. For yeast, we used the prior network created in [17]. Briefly, this approach scanned the promoter of genes (defined as 1,000 bp upstream of the first ATG of a gene) using TestMotif [47] using position weight matrices (PWMs) from Gordan et al. [48] and YeTFaSCo [49]. The edges in the prior network were weighted using percentile ranking where lowest p-value is assigned the highest rank. For mammalian prior networks, we used PIQ [50] which scores motif instances using DNase I footprints. Motif instances were mapped to ±10Kbp of the transcription start site (TSS). For mESC and mDC cell lines PWMs from the CISBP database [51] were used, while for hESC cell line, in addition to CIS-BP, PWMs from the ENCODE [52] and JASPAR [53] databases were also used. For estimation of footprints, DNase I assays from ENCODE were used [54]. To add an edge between a TF and a gene, we required that the TF’s accessible motif occur in the gene’s promoter. The edge weight between a TF and a target was the p-value of the motif instance overlapping the promoter; if there were multiple instances, the one with the most significant instance was used as the score. A threshold was applied to the score of the motif instances to create a network with similar edge density as the yeast motif prior network followed by percentile ranking on the edge score to create the final network. Inferelator has the option to estimate TFA internally using the input prior network. We additionally applied the original NCA algorithm [33], which uses an iterative approach to estimate TFA. Briefly, given gene expression matrix *E* and adjacency matrix *A*, NCA estimates the TFA matrix *P* in a way that 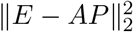. The iterative approach works by estimating P matrix from the given A matrix, and then updating the A matrix by fixing the P matrix. We evaluated the inferred networks using F-score, AUPR and predictable TFs of these algorithms and compared to the baseline versions that did not use priors and/or TFA and also correlation.

### Application of network inference algorithms

#### Execution and post-processing

Each algorithm was applied to the processed expression matrices on a local high-performance computing cluster using HTCondor https://chtc.cs.wisc.edu. Where possible, we deployed each algorithm in a stability selection mode. Specifically, we selected 50% of the cells from each expression matrix randomly, applied the algorithm to this subset, and repeated this process 100 times, averaging the interaction scores over the 100 subset networks to obtain a final inferred network. As some algorithms perform a similar downsampling internally (SCENIC, Inferelator), we did not run these algorithms within our external stability selection framework. Other algorithms could not be run 100 times due to excessive runtimes (PIDC, Scribe). After completing the runs of each algorithm, we formatted the variety of network formats to a single three-column file (transcription factor, target, score). The score varied depending upon the algorithm and could be a pairwise correlation (LEAP, Correlation), information theoretic dependence (knnDREMI, SCRIBE, PIDC), regression weight (SILGGM, SCODE), confidence (SCENIC, Inferelator). The magnitude of the score was interpreted as the strength of the interaction. We restricted all networks to include only edges where the “regulator” was a known TF from one of our experimentally derived gold standard networks. The final networks were used in the evaluations of network accuracy and similarity (detailed below) and are available in **Additional File 3**. The scripts for postprocessing the results are available at https://github.com/Roy-lab/scRNAseq_NetInference.

#### Estimation of pseudo time

Some algorithms (LEAP, SCODE, Scribe) incorporate knowledge of cellular trajectory and pseudotime, which was learned using Monocle 2 [55]. To avoid biasing the trajectories with a choice of marker genes, we followed the best practices in the Monocle documentation to learn trajectories in an unsupervised approach. In brief, we clustered the cells using Monocle’s density-peak clustering algorithm after dimensionality reduction with t-SNE. We next tested for genes differentially expressed across these clusters, and selected the top thousand most significantly differential genes (as measured by q-value) to use as marker genes. Finally, we learned a DDRTree embedding of the expression in each cell and applied Monocle’s cell ordering function to infer the position of each cell on a trajectory and their corresponding pseudotime assignments. We used Monocle trajectories for all datasets except the yeast Jackson dataset for which we used Diffusion Pseudo Time (DPT) [56] due to high runtime of the Monocle software.

### Description of gold standards

We curated multiple experimentally derived networks of regulatory interactions from published databases and the literature (**Additional File 3**) to serve as gold standards for our network inference algorithms. These experiments are typically based on ChIP-chip, ChIP-seq or regulator perturbation followed by global transcriptome profiling. We obtained multiple networks based on ChIP and TF perturbation experiments for each organism and cell type. When multiple ChIP or perturbation interactions were available we took a union of the networks. We refer to the ChIP derived gold standard as “ChIP” and the perturbation derived gold standard as “Perturb”. Finally, we took the intersection of these two unions, as the third primary gold standard networks for our evaluations of network accuracy (ChIP+Perturb).

Finally, we created a fourth embryonic stem cell (ESC) specific gold standard network from the primary literature [24–30] by conducting a literature survey of gene regulatory networks for the ESC state. Regulatory edges from the publications were manually extracted from network figures and further curated by a stem cell biologist (see **Acknowledgements**). The specific publications and figures used to create our curated gold standard are in **Additional File 2**, **Table S5**.

### Evaluation metrics

To evaluate the accuracy of the inferred networks, we used three metrics - Area under the Precision Recall curve (AUPR), F-score, and predictable TFs - between each network and each relevant gold standard network. Prior to computing each accuracy measure, we filtered both the inferred network and the gold standard to include only edges that contained TFs and target genes from the intersection of the gold standard’s gene set with the original expression matrix’s gene set. This was to eliminate any penalty for missing edges included in the gold standard that could not be inferred by the algorithm due to lack of expression in the original data.

#### Area Under the Precision Recall curve (AUPR)

We used AUPR to measure the accuracy of the global network structure. Specifically, we sorted the inferred edges in descending order of confidence, and computed the precision and recall with respect to a gold standard as edges were sequentially included. We report the area under this curve as AUPR. We used the script auc.java from https://github.com/Arun-George-Zachariah/AUC [57].

#### F-score

We used F-score of the top *x* edges, *x* ∈ {100,300, 500,1000,3000, 5000, 10000,30000, 50000} to measure the accuracy of the most confident edges in an inferred network. Specifically, the F-score is computed as the harmonic mean of the precision and recall of the top x edges with respect to a gold standard.

#### Predictable TFs

We used predictable TFs to obtain a granular measure of network accuracy at the individual TF’s target set. Specifically, for each TF in the inferred network, we use the hypergeometric test to assess whether the overlap between the predicted targets and the targets in the gold standard was significantly higher than random. The P-values were corrected using FDR for multiple hypothesis correction. We considered a P-value < 0.05 as significant overlap. Each algorithm was ranked based on the number of predictable TFs.

## Availability of data and materials

Datasets used in our experiments and analyses have been deposited with DOI `10.5281/zenodo.5909090`, along with the following groups of files which were too large to include in the manuscript as supplementary materials:

- Gold standard networks used as ground truth to measure accuracy of the inferred networks.
- Networks generated from network inference methods.

All scripts for executing and evaluating the algorithms are available at https://github.com/Roy-lab/scRNAseq_NetInference.

## Author’s contributions

M.S. and S.R. conceptualized the project. M.S., S.G.M, A.F.S, V.P. performed data download and preprocessing. M.S., S.G.M., A.F.S, V.P., J.L., S.P. performed benchmarking experiments. M.S., A.F.S., J.L, S.P., and J.S. ran evaluation scripts for examining results. J.L., S.G.M, M.S., S.P., and S.R. wrote the paper.

## Acknowledgements

This work was supported by the NIH grants R01GM117339 and U01EB029371, a UW Data Science Foundation grant. M.S. acknowledges support from an NIH T15 training fellowship (T15 LM007359), S.G.M acknowledges support from an NIH T32 training fellowship (T32 GM007133), and S.P. acknowledges support from DOE grant DE-SC0021052 and NSF grant 2010789. The authors thank the Center for High-throughput Computing for enabling the large-scale computational experiments of this project. We would like to thank Dr. Rupa Sridharan for her guidance in creation of the literature curated gold standard network for the embryonic stem cell state.

## Additional Files

**Additional file 1 — Supplementary Tables**

This file has all the supplementary tables, S1-S5. **Table S1** has the summary of algorithms used in comparisons and benchmarking. **Table S2** has the summary and statistics of datasets used for algorithm comparisons listing reference, GEO ID, species, cell type, number of cells before and after filtering was applied, number of genes before and after filtering was applied, and brief dataset description.**Table S3** has statistics of our gold standard networks including the number of regulators, targets and edges. It also indicates which gold standards are used for which dataset. **Table S4** has the description and data source of bulk RNA-seq datasets. **Table S5** provides the list of publications and figures used to make the mESC literature curated gold standard.

**Additional file 2 — Supplementary Datasets**

Expression datasets, regulator sets and gold standard networks for each of the biological conditions considered.

**Additional file 3 — Supplemental Figures**

All supplemental figures are documented in this pdf file.

**Additional file 4 — Supplementary Text**

This text describes the effects of dataset size and depth on the performance of the network inference methods.

