## Supplemental Data 1 for "Identifying strengths and weaknesses of methods for computational network inference from single cell RNA-seq data"

### Supplementary figures

S11 Network inference utilizing gene expression alone compared to addition of priors and TFA . 23

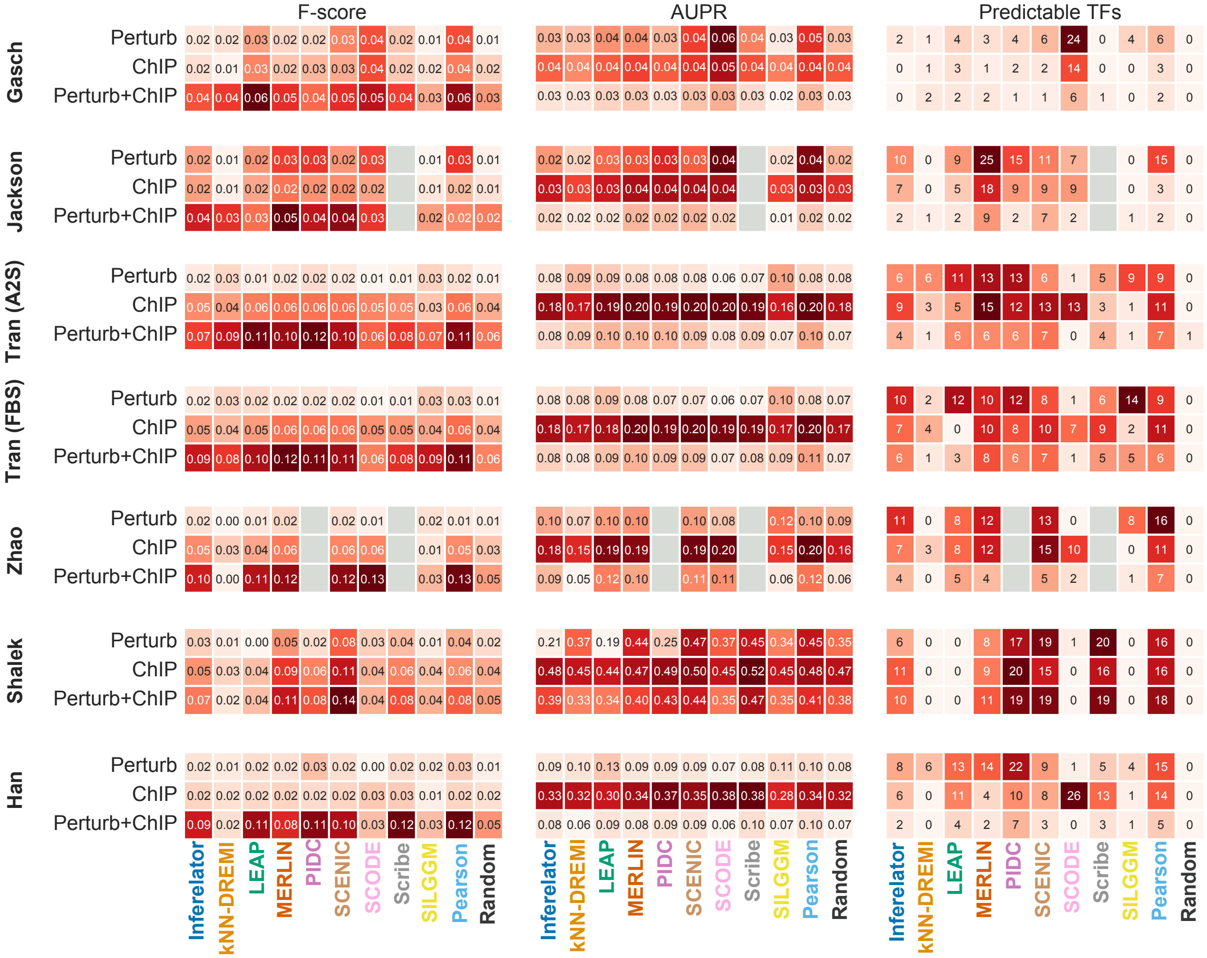
F-score
AUPR
Predictable TFs
Gasch
Jackson
Tran (A2S)
Tran (FBS)
Zhao
Shalek
Han
Perturb
ChIP
Perturb+ChIP
Inferelator
kNN-DREMI
LEAP
MERLIN
PIDC
SCENIC
SCODE
Scribe
SILGGM
Pearson
Random

**Figure S1.** Evaluation metrics for network inference on individual datasets. Heatmaps depict the values of each metric used for evaluating inferred networks. From left to right: F-score of top 5,000 edges, Area under the precision recall (AUPR) curve, and the number of predictable TFs. Algorithms are ordered alphabetically, followed by the Pearson and random networks.

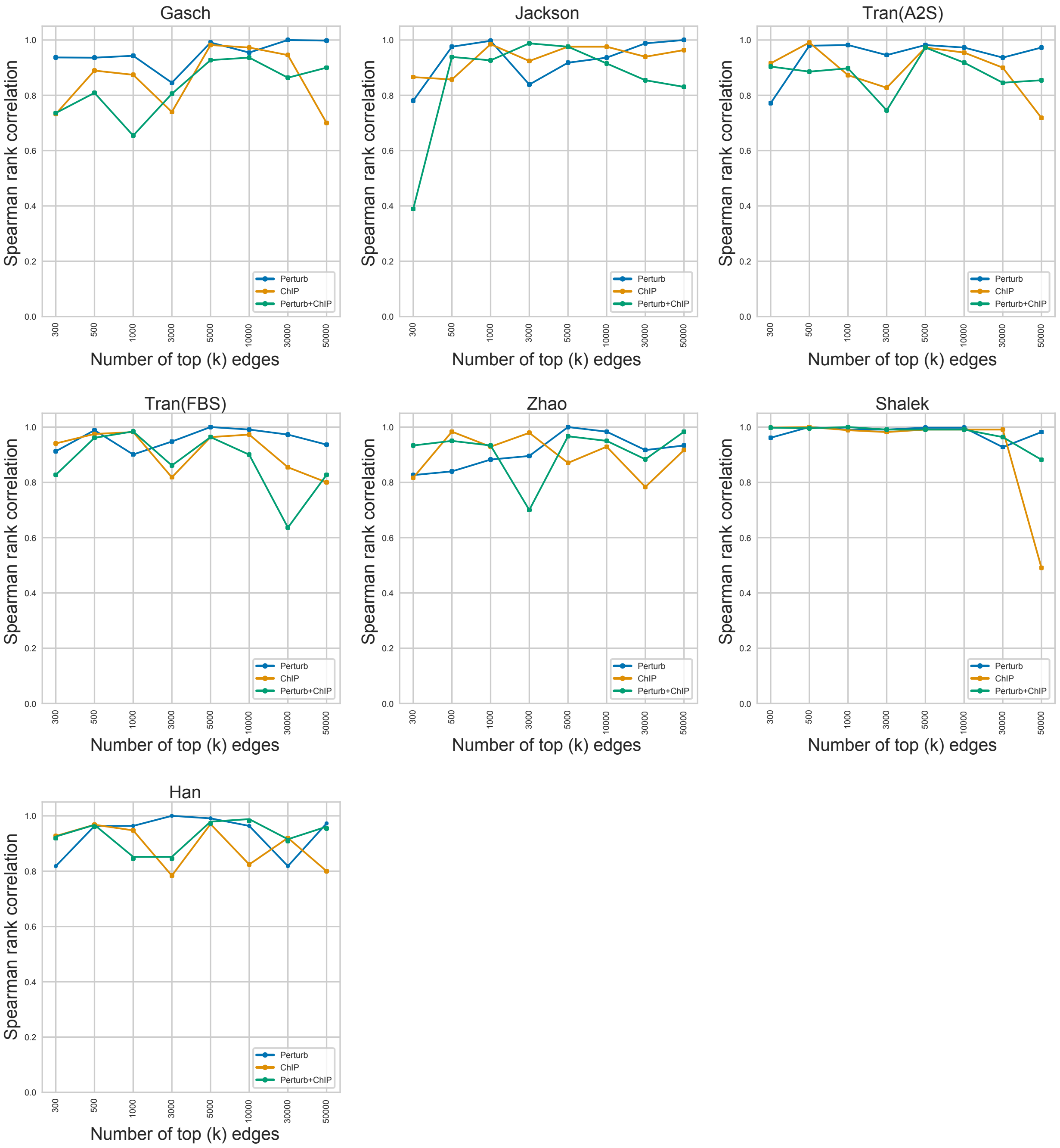
Gasch
Jackson
Tran(A2S)
Tran(FBS)
Zhao
Shalek
Han
Spearman rank correlation
Number of top (k) edges
0.0
0.2
0.4
0.6
0.8
1.0
300
500
1000
3000
5000
10000
30000
50000
Perturb
ChIP
Perturb+ChIP

**Figure S2.** Effect of number of top edges selected on F-score based algorithm rankings. We measured the F-score of each algorithm with respect to each experimentally derived network using the top 100, 300, 500, 1,000, 3,000, 5,000, 10,000, 30,000, and 50,000 edges. We computed the Spearman rank correlation between algorithm performances for each consecutive pair of edge sets, that is, the value corresponding to 300 on each graph shows the correlation between algorithm ranks when using the top 100 and top 300 edges. We found that across datasets, relative algorithm performance generally stabilized when using the top 5,000 edges and used this threshold for our analyses.

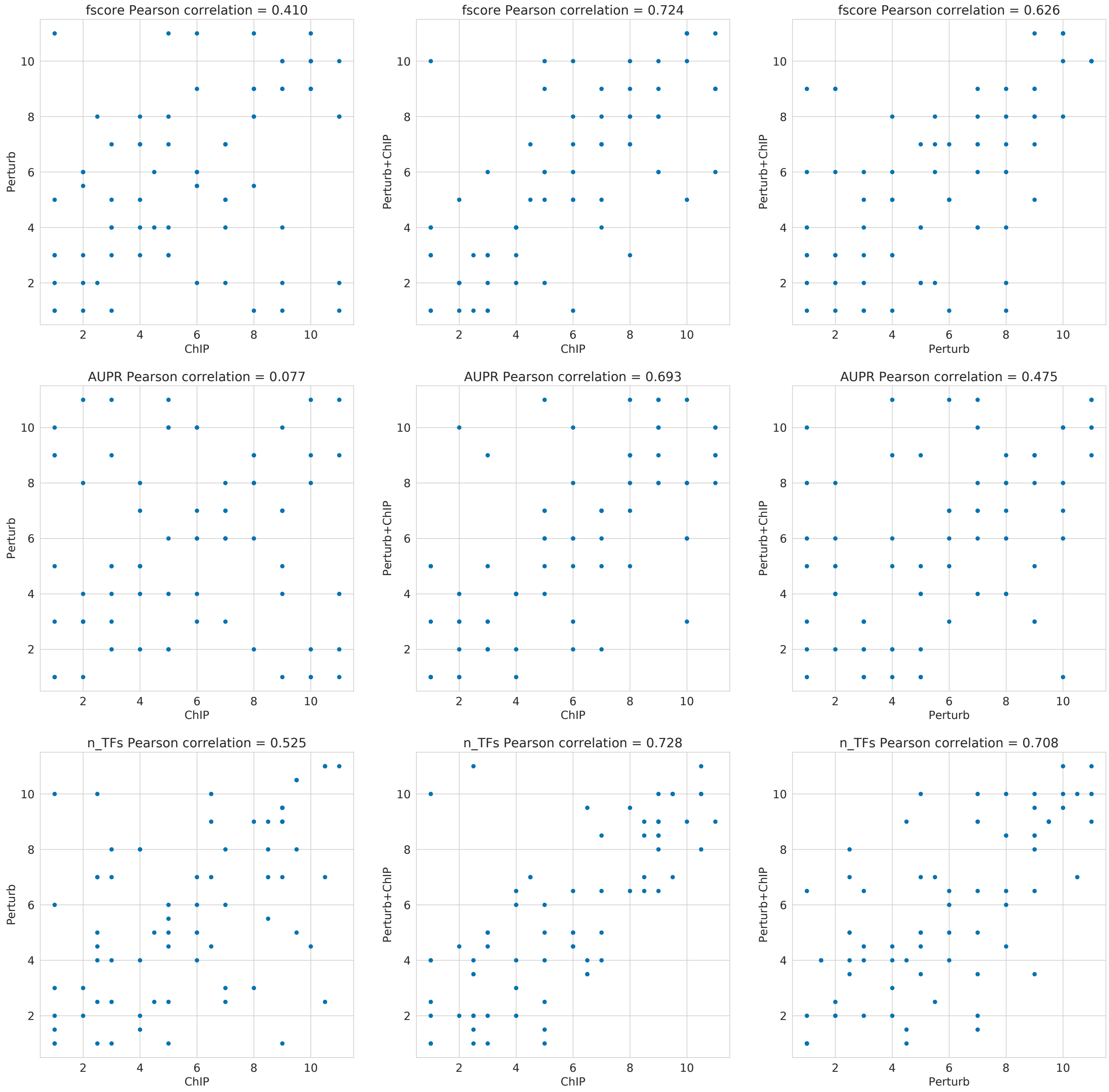
fscore Pearson correlation = 0.410
fscore Pearson correlation = 0.724
fscore Pearson correlation = 0.626
AUPR Pearson correlation = 0.077
AUPR Pearson correlation = 0.693
AUPR Pearson correlation = 0.475
n_TFs Pearson correlation = 0.525
n_TFs Pearson correlation = 0.728
n_TFs Pearson correlation = 0.708
ChIP
Perturb
Perturb+ChIP

**Figure S3.** Pearson correlation coefficients of ranks between gold standards when using different network inference evaluation metrics. Scatter plots show the correlations of algorithm ranks between pairs of gold standards (Perturb, ChIP, and Perturb+ChIP). The Pearson correlation coefficients are calculated based on ranks on the three evaluation metrics: AUPR, F-score, predictable TFs (n_TFs).

#### ChIP

| | SCENIC | Pearson | MERLIN | PIDC | SCODE | LEAP | Scribe | Inferelator | kNN-DREMI | SILGGM |
|---|---|---|---|---|---|---|---|---|---|---|
| Number of predictable TFs | 10 | 11 | 10 | 8 | 7 | 0 | 9 | 7 | 4 | 2 |
| Sox2 | 1.6 | 1.6 | 1.6 | 1.6 | 1.3 | | 1.4 | 1.5 | | |
| Esrrb | 1.4 | 1.4 | 1.3 | 1.4 | | | | 1.6 | | 1.2 |
| Nanog | 1.5 | 1.4 | 1.3 | 1.4 | | | 1.2 | 1.4 | | |
| Sall4 | 1.7 | 1.9 | 1.8 | 2.2 | | | 1.8 | 1.9 | | |
| Nr0b1 | 2.8 | 2.7 | 2.1 | 2.4 | | | | 3.1 | | 1.5 |
| Pou5f1 | 1.3 | 1.4 | 1.2 | 1.4 | | | 1.4 | | | |
| Klf2 | 3.3 | 8.5 | | 8.1 | | | 2.5 | 2.7 | 10.5 | |
| E2f4 | | 1.4 | 1.5 | | 1.5 | | | 1.5 | | |
| Jarid2 | | | | | | | | | 2.6 | |
| Mycn | 1.4 | 1.3 | | | | | 1.3 | | | |
| Zic3 | | 3.1 | | 3.1 | | | 5.1 | | | |
| Myc | 1.3 | | 1.3 | | 1.3 | | | | | |
| Zic1 | | | | | | | | | | |
| Suz12 | | | | | | | | | 2.7 | |
| Fosl2 | | | | | | | | | | |
| Gbx2 | | | | | | | | | | |
| Foxc1 | | | | | | | | | | |
| Zfx | | | 1.8 | | 1.5 | | | | | |
| Chd1 | 2.8 | | | | 2.2 | | | | | |
| Dmrt1 | | | | | | | | | | |
| Id1 | | | | | | | | | | |
| Jun | | | | | | | | | | |
| Sox15 | | | | | | | | | | |
| Stat3 | | | | | 1.6 | | | | | |
| Phc1 | | | | | | | | | 2.4 | |
| Tbx3 | | | 2.8 | | | | | | | |
| Zfp42 | | | | | | | 1.6 | | | |
| Klf4 | | | | | 1.5 | | | | | |
| Otx2 | | | | | | | | | | |
| Trim28 | | | | | | | 1.2 | | | |
| Klf5 | | 11.8 | | | | | | | | |
| Lhx2 | | | | | | | | | | |
| Smad7 | | | | | | | | | | |
| Gata2 | | | | | | | | | | |
| Etv3 | | | | | | | | | | |
| Mybl2 | | | | | | | | | | |

#### Perturb

| | SCENIC | Pearson | MERLIN | PIDC | SCODE | LEAP | Scribe | Inferelator | kNN-DREMI | SILGGM |
|---|---|---|---|---|---|---|---|---|---|---|
| Number of predictable TFs | 8 | 9 | 10 | 12 | 1 | 12 | 6 | 10 | 2 | 14 |
| Sox2 | 1.5 | 1.4 | 1.5 | 1.6 | | 1.7 | 1.3 | 1.7 | | 1.7 |
| Esrrb | 3.0 | 2.2 | 2.9 | 2.4 | | 1.4 | 3.4 | 3.5 | | 2.6 |
| Nanog | 2.1 | 2.1 | 2.0 | 2.1 | | 2.1 | 1.7 | 2.4 | | 2.3 |
| Sall4 | | 1.6 | 1.5 | 1.9 | | 1.7 | 1.5 | 1.7 | 2.0 | 1.8 |
| Nr0b1 | 4.7 | | 3.9 | | | 1.7 | | 5.3 | | 3.6 |
| Pou5f1 | | 1.7 | 1.7 | 1.6 | | 1.4 | | | | 1.3 |
| Klf2 | | | | | | | | | | |
| E2f4 | | | | | | | | | | |
| Jarid2 | 2.4 | 3.0 | | 2.7 | | | 1.9 | 2.7 | | |
| Mycn | | | | | | | | | | |
| Zic3 | | | | | | | | | | |
| Myc | | | | | | | | | | |
| Zic1 | 4.3 | | 4.0 | 12.2 | | | | 6.8 | | 2.6 |
| Suz12 | | | | | 3.5 | | 4.6 | | | |
| Fosl2 | | 2.5 | | 2.3 | | 1.9 | | 1.7 | | |
| Gbx2 | | 1.9 | | 2.0 | | 2.3 | | | | 1.6 |
| Foxc1 | 2.2 | | 2.0 | 3.4 | | | | 2.4 | | |
| Zfx | | | | | | | | | | |
| Chd1 | | | | | | | | | | |
| Dmrt1 | | 1.7 | 1.5 | | | | | | | 1.2 |
| Id1 | | | | 8.6 | | 22.9 | | 4.8 | | |
| Jun | 2.3 | | | 2.8 | | 2.1 | | | | |
| Sox15 | | | | | | 2.2 | | | | 2.3 |
| Stat3 | | | | | | | | | | |
| Phc1 | | | | | | | | | | |
| Tbx3 | | | | | | | | | | |
| Zfp42 | | | | | | | | | | |
| Klf4 | | | | | | | | | | |
| Otx2 | | | 2.6 | | | | | | | 2.0 |
| Trim28 | | | | | | | | | | |
| Klf5 | | | | | | | | | | |
| Lhx2 | | | | | | | | | | 2.2 |
| Smad7 | | | | | | 1.6 | | | | |
| Gata2 | | | | | | | | | 1.4 | |
| Etv3 | | | | | | | | | | 10.2 |
| Mybl2 | | | | | | | | | | 2.6 |

Number of predictable TFs

| | | | | | |
|---|---|---|---|---|---|
| 0.0 | 4.0 | 8.0 | 12.0 | 16.0 | 20.0 |

Fold enrichment

| | | | | | |
|---|---|---|---|---|---|
| 0.0 | 1.0 | 2.0 | 3.0 | 4.0 | 5.0 |

**Figure S4.** Predictable TFs for the Tran (FBS) dataset. Heatmaps show the enrichment of each transcription factor's target set from a perturbation-based (left) or ChIP-based (right) experimentally derived gold standard network in an inferred network. Columns are ordered based on overall ranking of methods. Individual white cells indicate TFs that were not considered as a predictable TF by a method. An entire row of white cells indicates the TF did not appear in one of the two experimentally derived networks.

#### ChIP

| | SCENIC | Pearson | MERLIN | PIDC | SCODE | LEAP | Scribe | Inferelator | kNN-DREMI | SILGGM |
|---|---|---|---|---|---|---|---|---|---|---|
| Number of predictable TFs | 15 | 11 | 12 | 0 | 10 | 8 | 0 | 7 | 3 | 0 |
| Nanog | 1.3 | 1.3 | 1.2 | | 1.3 | 2.0 | | 1.3 | | |
| Pou5f1 | 1.1 | 1.2 | 1.2 | | 1.2 | 1.1 | | | | |
| Sox2 | 2.0 | 1.7 | 1.5 | | 1.6 | | | 1.6 | | |
| Esrrb | 1.4 | 1.4 | 1.3 | | | 1.3 | | | | |
| Cdx2 | 2.1 | 4.5 | 1.6 | | | | | | | |
| Sall4 | | 1.5 | 2.6 | | | 1.6 | | | | |
| Sox17 | 1.4 | | | | 1.5 | | | | | |
| E2f4 | | | 1.5 | | 1.5 | | | 1.4 | 1.2 | |
| Zfp42 | 1.6 | 1.4 | 1.6 | | 2.0 | | | | | |
| Klf4 | 1.5 | | | | 1.6 | 1.1 | | | | |
| Nr0b1 | 2.8 | 3.4 | 2.6 | | | | | | | |
| Klf2 | 5.4 | 4.9 | | | | | | 8.1 | | |
| Zfx | | | | | 2.1 | | | 1.5 | 1.3 | |
| Sox11 | | | | | | | | | | |
| Stat3 | 2.8 | | 2.4 | | | | | | | |
| Max | 1.6 | | | | 1.7 | | | | | |
| Tbx3 | 2.4 | | 1.9 | | | | | | | |
| Jarid2 | | 2.3 | | | | 2.1 | | | | |
| Gata3 | | | | | | | | | | |
| Chd1 | | | 3.0 | | | | | 2.7 | | |
| Gata2 | | | | | | | | | | |
| Myod1 | | | | | | | | | | |
| Atf3 | | | | | | | | | | |
| Otx2 | | | | | | | | | | |
| Nr2f2 | | | | | | | | | | |
| Dmrt1 | | | | | | | | | | |
| Tcf3 | | | | | | 1.4 | | | | |
| Chd7 | | | | | | | | 7.1 | | |
| Yy1 | 4.0 | | | | | | | | | |
| Mycn | | | | | 2.1 | | | | | |
| Suz12 | | | | | | | | | 1.4 | |
| Trim28 | | | | | | 1.7 | | | | |
| Prdm14 | | 2.1 | | | | | | | | |
| Klf5 | | | | | | | | | | |
| Myc | 2.2 | | | | | | | | | |
| Etv1 | | | | | | | | | | |
| Foxa1 | | | | | | | | | | |
| Tcea3 | | | | | | | | | | |
| T | | | | | | | | | | |
| Nr2f1 | | | | | | | | | | |
| Id1 | | | | | | | | | | |
| Otx1 | | | | | | | | | | |
| Etv5 | | | | | | | | | | |
| Ascl1 | | | | | | | | | | |
| Gbx2 | | | | | | | | | | |
| Jun | | | | | | | | | | |
| Mef2c | | | | | | | | | | |

#### Perturb

| | SCENIC | Pearson | MERLIN | PIDC | SCODE | LEAP | Scribe | Inferelator | kNN-DREMI | SILGGM |
|---|---|---|---|---|---|---|---|---|---|---|
| Number of predictable TFs | 13 | 16 | 12 | 0 | 0 | 8 | 0 | 11 | 0 | 8 |
| Nanog | 2.0 | 1.7 | 1.6 | | | | | 1.8 | | |
| Pou5f1 | 1.3 | 1.7 | 1.3 | | | 1.5 | | 1.4 | | |
| Sox2 | 1.7 | 1.5 | 1.8 | | | | | 1.6 | | |
| Esrrb | 3.0 | 3.0 | | | | | | 2.7 | | |
| Cdx2 | 1.4 | | 1.4 | | | | | 1.5 | | |
| Sall4 | | 2.6 | | | | 1.9 | | | | |
| Sox17 | 2.6 | 2.8 | | | | 2.0 | | 2.4 | | |
| E2f4 | | | | | | | | | | |
| Zfp42 | | | | | | | | | | |
| Klf4 | | 1.4 | | | | 1.4 | | | | |
| Nr0b1 | | 5.7 | | | | | | | | |
| Klf2 | | | | | | | | | | |
| Zfx | | | | | | | | | | |
| Sox11 | 2.4 | 1.6 | 2.8 | | | 1.6 | | 1.9 | | |
| Stat3 | | | | | | | | | | |
| Max | | | | | | | | | | |
| Tbx3 | | | | | | | | | | |
| Jarid2 | | | | | | | | | | |
| Gata3 | 1.6 | | 1.5 | | | 2.8 | | 1.9 | | |
| Chd1 | | | | | | | | | | |
| Gata2 | 1.6 | | 1.6 | | | | | 1.6 | | |
| Myod1 | 3.0 | | 2.5 | | | | | | | 2.2 |
| Atf3 | 5.9 | | 4.6 | | | | | 10.2 | | |
| Otx2 | | 2.3 | 4.7 | | | | | | | 2.3 |
| Nr2f2 | 2.3 | 1.7 | | | | 1.7 | | | | |
| Dmrt1 | | 1.8 | 1.6 | | | | | | | |
| Tcf3 | | | | | | | | | | |
| Chd7 | | | | | | | | | | |
| Yy1 | | | | | | | | | | |
| Mycn | | | | | | | | | | |
| Suz12 | | | | | | | | | | |
| Trim28 | | | | | | | | | | |
| Prdm14 | | | | | | | | | | |
| Klf5 | | 4.9 | | | | 5.3 | | | | |
| Myc | | | | | | | | | | |
| Etv1 | | 8.0 | | | | | | | | |
| Foxa1 | | | | | | | | | | 1.3 |
| Tcea3 | | | | | | | | | | 4.6 |
| T | | | | | | | | | | 1.7 |
| Nr2f1 | | 4.6 | | | | | | | | |
| Id1 | | | 4.8 | | | | | | | |
| Otx1 | | | | | | | | | | 2.1 |
| Etv5 | | | | | | | | 4.6 | | |
| Ascl1 | | | | | | | | | | 1.7 |
| Gbx2 | 2.0 | | | | | | | | | |
| Jun | | 3.3 | | | | | | | | |
| Mef2c | | | | | | | | | | 2.5 |

Number of predictable TFs

| | | | | | |
|---|---|---|---|---|---|
| 0.0 | 4.0 | 8.0 | 12.0 | 16.0 | 20.0 |

Fold enrichment

| | | | | | |
|---|---|---|---|---|---|
| 0.0 | 1.0 | 2.0 | 3.0 | 4.0 | 5.0 |

**Figure S5.** Predictable TFs for the Zhao dataset. Heatmaps show the enrichment of each transcription factor's target set from a perturbation-based (left) or ChIP-based (right) experimentally derived network in an inferred network. Columns are ordered based on overall ranking of methods. Individual white cells indicate transcription factors that were not considered as a predictable TF by a method. An entire row of white cells indicates the TF did not appear in one of the two experimentally derived networks.

#### ChIP

| | SCENIC | Pearson | MERLIN | PIDC | SCODE | LEAP | Scribe | Inferelator | kNN-DREMI | SILGGM |
|---|---|---|---|---|---|---|---|---|---|---|
| Number of predictable TFs | 15 | 16 | 9 | 20 | 0 | 0 | 16 | 11 | 0 | 0 |
| Atf3 | 1.3 | 1.3 | 1.2 | 1.4 | | | 1.3 | 1.2 | | |
| Rel | 1.5 | 1.5 | 1.3 | 1.8 | | | 1.5 | 1.5 | | |
| Stat1 | 1.1 | 1.1 | 1.2 | 1.2 | | | 1.1 | 1.1 | | |
| Nfkb1 | 2.0 | 2.0 | 1.8 | 2.3 | | | 1.5 | 2.0 | | |
| Stat3 | 1.1 | 1.2 | 1.2 | 1.3 | | | 1.4 | 1.2 | | |
| Relb | 1.5 | 1.5 | 1.4 | 1.7 | | | 1.6 | 1.4 | | |
| Stat2 | 1.1 | 1.1 | | 1.2 | | | 1.2 | 1.1 | | |
| Egr2 | 1.2 | 1.2 | | 1.2 | | | 1.2 | 1.2 | | |
| Cebpb | 1.1 | 1.1 | | 1.0 | | | 1.1 | | | |
| Irf1 | 1.1 | 1.1 | | 1.2 | | | 1.1 | | | |
| Junb | 1.1 | 1.1 | 1.1 | 1.2 | | | 1.1 | | | |
| Egr1 | 1.3 | 1.3 | | 1.3 | | | | 1.2 | | |
| Maff | 1.5 | 1.4 | 1.4 | 1.6 | | | | | | |
| Irf2 | 2.3 | 3.5 | 2.5 | 2.9 | | | | 2.0 | | |
| Ets2 | 1.2 | | | 1.1 | | | 1.2 | | | |
| Irf4 | | 1.1 | | 1.1 | | | 1.3 | | | |
| Rela | | | | 1.2 | | | 1.1 | | | |
| Jak2 | | | | | | | | | | |
| Runx1 | | 1.3 | | 1.3 | | | | | | |
| Hif1a | | | | 1.6 | | | 1.6 | | | |
| Atf4 | | | | 1.4 | | | | 1.3 | | |
| E2f1 | | | | | | | 1.3 | | | |
| Hsp90b1 | | | | | | | | | | |
| Ctcf | | | | | | | | | | |
| Nek6 | | | | | | | | | | |
| Actb | | | | | | | | | | |
| Ahr | | | | | | | | | | |
| Ly96 | | | | | | | | | | |

#### Perturb

| | SCENIC | Pearson | MERLIN | PIDC | SCODE | LEAP | Scribe | Inferelator | kNN-DREMI | SILGGM |
|---|---|---|---|---|---|---|---|---|---|---|
| Number of predictable TFs | 19 | 16 | 8 | 17 | 1 | 0 | 20 | 6 | 0 | 0 |
| Atf3 | 1.4 | 1.3 | 1.2 | 1.6 | | | 1.5 | 1.2 | | |
| Rel | 1.1 | 1.1 | 1.1 | 1.1 | | | 1.1 | | | |
| Stat1 | 1.1 | 1.1 | | 1.1 | | | 1.1 | 1.1 | | |
| Nfkb1 | 1.1 | 1.1 | 1.1 | 1.1 | | | | 1.1 | | |
| Stat3 | 1.1 | 1.1 | | 1.2 | | | 1.1 | | | |
| Relb | 1.3 | 1.1 | | 1.2 | | | 1.3 | | | |
| Stat2 | 1.1 | 1.1 | 1.1 | 1.1 | | | 1.1 | | | |
| Egr2 | 1.2 | 1.1 | | 1.3 | | | 1.3 | | | |
| Cebpb | 1.6 | 1.6 | 1.4 | 1.5 | | | 1.6 | 1.4 | | |
| Irf1 | 1.3 | 1.3 | 1.2 | 1.4 | | | 1.3 | | | |
| Junb | 1.4 | | | | | | 1.5 | | | |
| Egr1 | 1.7 | 1.3 | | 1.7 | | | 2.2 | | | |
| Maff | 1.2 | 1.1 | 1.1 | | | | | | | |
| Irf2 | | | | | | | 1.3 | | | |
| Ets2 | 1.3 | 1.3 | | 1.5 | | | 1.3 | | | |
| Irf4 | 1.3 | 1.1 | | 1.4 | | | 1.4 | | | |
| Rela | 1.1 | 1.1 | | | | | 1.2 | | | |
| Jak2 | 2.2 | 2.1 | 1.8 | 2.6 | | | 1.9 | 2.9 | | |
| Runx1 | | | | 1.1 | | | 1.2 | | | |
| Hif1a | 1.2 | | | | | | 1.5 | | | |
| Atf4 | | | | | | | | | | |
| E2f1 | | | | | | | 1.6 | | | |
| Hsp90b1 | | | | | | | | 13.2 | | |
| Ctcf | | | | | | | 1.5 | | | |
| Nek6 | | | | | 34.7 | | | | | |
| Actb | | | | 3.5 | | | | | | |
| Ahr | 1.3 | | | | | | | | | |
| Ly96 | | | | 11.6 | | | | | | |

Number of predictable TFs

| | | | | | |
|---|---|---|---|---|---|
| 0.0 | 4.0 | 8.0 | 12.0 | 16.0 | 20.0 |

Fold enrichment

| | | | | | |
|---|---|---|---|---|---|
| 0.0 | 1.0 | 2.0 | 3.0 | 4.0 | 5.0 |

**Figure S6.** Predictable TFs for the Shalek dataset. Heatmaps show the enrichment of each transcription factor's target set from a perturbation-based (left) or ChIP-based (right) experimentally derived network in an inferred network. Columns are ordered based on overall ranking of methods. Individual white cells indicate transcription factors that were not considered as a predictable TF by a method. An entire row of white cells indicates the TF did not appear in one of the two experimentally derived networks.

#### ChIP

| | SCENIC | Pearson | MERLIN | PIDC | SCODE | LEAP | Scribe | Inferelator | kNN-DREMI | SILGGM |
|---|---|---|---|---|---|---|---|---|---|---|
| Number of predictable TFs | 2 | 3 | 1 | 2 | 14 | 3 | 0 | 0 | 1 | 0 |
| MSN2 | 2.8 | 1.9 | | 2.5 | 4.4 | 2.0 | | | 2.7 | |
| IFH1 | 2.9 | 7.7 | 4.0 | | | 5.0 | | | | |
| SFP1 | | 4.7 | | | 3.3 | 4.8 | | | | |
| XBP1 | | | | | | | | | | |
| CIN5 | | | | 1.9 | 3.8 | | | | | |
| GIS1 | | | | | | | | | | |
| GCN4 | | | | | | | | | | |
| YAP1 | | | | | | | | | | |
| GCR1 | | | | | 15.9 | | | | | |
| HSF1 | | | | | 4.2 | | | | | |
| GCR2 | | | | | 6.5 | | | | | |
| RAP1 | | | | | 2.3 | | | | | |
| TYE7 | | | | | 9.9 | | | | | |
| STE12 | | | | | 2.5 | | | | | |
| SOK2 | | | | | 2.7 | | | | | |
| PDR1 | | | | | 3.0 | | | | | |
| PHD1 | | | | | 2.1 | | | | | |
| RPN4 | | | | | | | | | | |
| HFI1 | | | | | | | | | | |
| FHL1 | | | | | 3.2 | | | | | |
| SKO1 | | | | | 3.1 | | | | | |
| SWI3 | | | | | | | | | | |
| RSF2 | | | | | | | | | | |
| MOT3 | | | | | | | | | | |
| AFT1 | | | | | | | | | | |
| TUP1 | | | | | | | | | | |
| MBF1 | | | | | | | | | | |
| SIN3 | | | | | | | | | | |
| REB1 | | | | | | | | | | |
| SNF6 | | | | | | | | | | |
| SNF2 | | | | | | | | | | |
| LEU3 | | | | | | | | | | |
| ABF1 | | | | | | | | | | |
| TEC1 | | | | | | | | | | |
| CUP2 | | | | | | | | | | |
| MGA2 | | | | | | | | | | |
| CST6 | | | | | | | | | | |
| SPT4 | | | | | | | | | | |
| HOG1 | | | | | | | | | | |

#### Perturb

| | SCENIC | Pearson | MERLIN | PIDC | SCODE | LEAP | Scribe | Inferelator | kNN-DREMI | SILGGM |
|---|---|---|---|---|---|---|---|---|---|---|
| Number of predictable TFs | 6 | 6 | 3 | 4 | 24 | 4 | 0 | 2 | 1 | 4 |
| MSN2 | 2.3 | 2.1 | 2.2 | 3.1 | 2.1 | 2.1 | | 2.1 | 3.5 | |
| IFH1 | | | | | | | | | | |
| SFP1 | | | | | 1.6 | | | | | |
| XBP1 | 4.5 | 3.9 | | 7.0 | | 4.8 | | 4.3 | | |
| CIN5 | | 6.2 | | | | | | | | |
| GIS1 | 4.1 | 3.9 | | 5.1 | 3.6 | | | | | |
| GCN4 | 1.7 | 1.7 | 1.8 | | | 1.6 | | | | |
| YAP1 | 2.0 | 2.4 | 2.1 | | 2.2 | | | | | |
| GCR1 | | | | | 3.6 | | | | | |
| HSF1 | | | | | 3.8 | | | | | |
| GCR2 | | | | | 5.7 | | | | | |
| RAP1 | | | | | 4.9 | | | | | |
| TYE7 | | | | | | | | | | 8.1 |
| STE12 | | | | | | | | | | |
| SOK2 | | | | | | | | | | |
| PDR1 | | | | | | | | | | |
| PHD1 | | | | | | | | | | |
| RPN4 | 2.6 | | | | 2.5 | | | | | |
| HFI1 | | | | | 6.0 | | | | | 4.8 |
| FHL1 | | | | | | | | | | |
| SKO1 | | | | | | | | | | |
| SWI3 | | | | | 3.8 | | | | | |
| RSF2 | | | | | 15.7 | | | | | |
| MOT3 | | | | | 51.6 | | | | | |
| AFT1 | | | | | 2.7 | | | | | |
| TUP1 | | | | | 4.1 | | | | | |
| MBF1 | | | | | | 5.3 | | | | |
| SIN3 | | | | | 5.7 | | | | | |
| REB1 | | | | | 5.8 | | | | | |
| SNF6 | | | | | 3.6 | | | | | |
| SNF2 | | | | | 3.9 | | | | | |
| LEU3 | | | | | 5.0 | | | | | |
| ABF1 | | | | | 3.0 | | | | | |
| TEC1 | | | | | | | | | | 7.0 |
| CUP2 | | | | 7.3 | | | | | | |
| MGA2 | | | | | 3.1 | | | | | |
| CST6 | | | | | 4.4 | | | | | |
| SPT4 | | | | | | | | | | 1.6 |
| HOG1 | | | | | 11.4 | | | | | |

Number of predictable TFs

| | | | | | |
|---|---|---|---|---|---|
| 0.0 | 4.0 | 8.0 | 12.0 | 16.0 | 20.0 |

Fold enrichment

| | | | | | |
|---|---|---|---|---|---|
| 0.0 | 1.0 | 2.0 | 3.0 | 4.0 | 5.0 |

**Figure S7.** Predictable TFs for the Gasch dataset. Heatmaps show the enrichment of each transcription factor's target set from a perturbation-based (left) or ChIP-based (right) experimentally derived network in an inferred network. Columns are ordered based on overall ranking of methods. Individual white cells indicate transcription factors that were not considered as a predictable TF by a method. An entire row of white cells indicates the TF did not appear in one of the two experimentally derived networks.

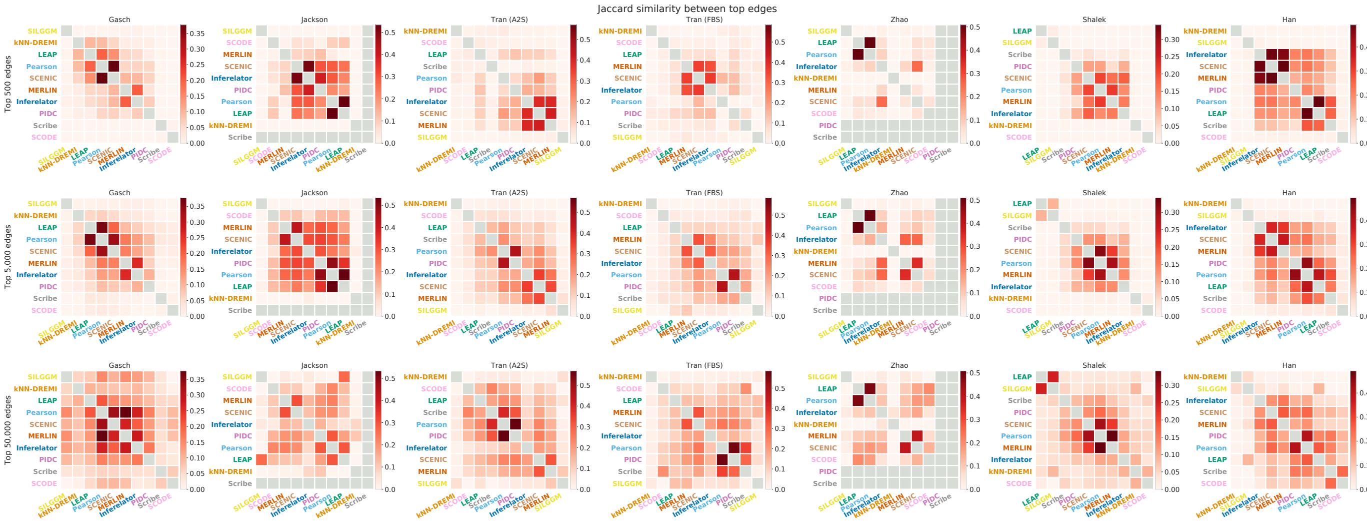
Jaccard similarity between top edges
Top 500 edges
Top 5,000 edges
Top 50,000 edges
Gasch
Jackson
Tran (A2S)
Tran (FBS)
Zhao
Shalek
Han
SILGGM
kNN-DREMI
LEAP
Pearson
SCENIC
MERLIN
Inferelator
PIDC
Scribe
SCODE

**Figure S8.** Jaccard similarity between top edge sets of inferred networks. Each heatmap showing inter-algorithm similarity between the networks inferred on each dataset (title), computed for the top 500, 5,000, and 50,000 edges. For each dataset, columns are ordered with respect to a hierarchical clustering of the similarity matrix based on the top 5,000 edges.

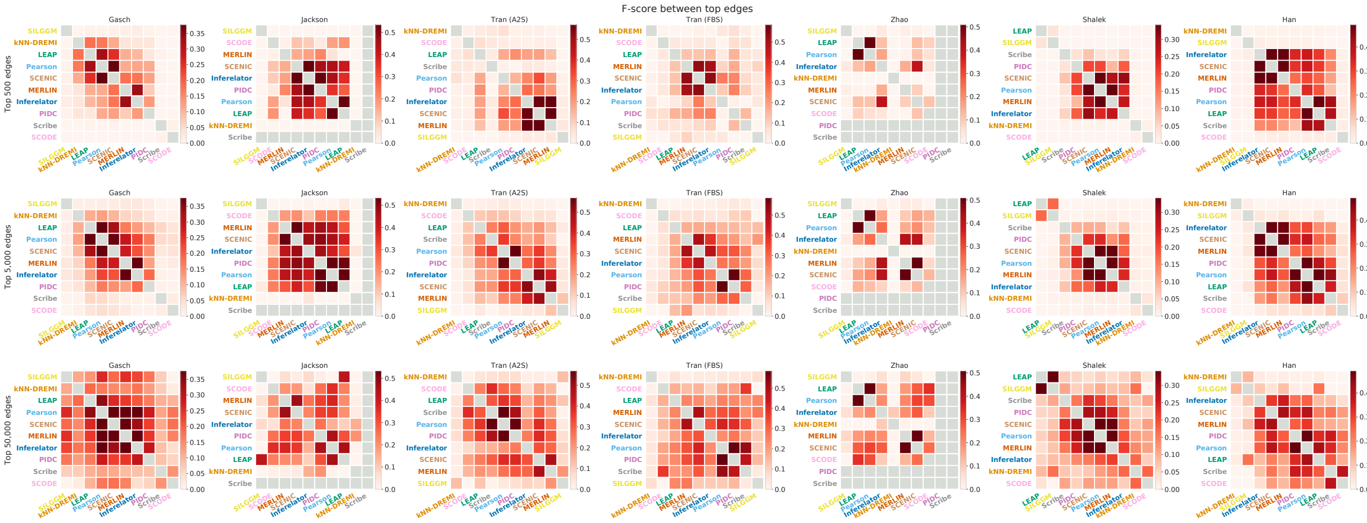

**Figure S9.** F-score similarity between top edge sets of inferred networks. Heatmaps showing inter-algorithm similarity between the networks inferred on each dataset, computed for the top 500, 5,000, and 50,000 edges. For each dataset, columns are ordered with respect to a hierarchical clustering of the F-score similarity matrix based on the top 5,000 edges.

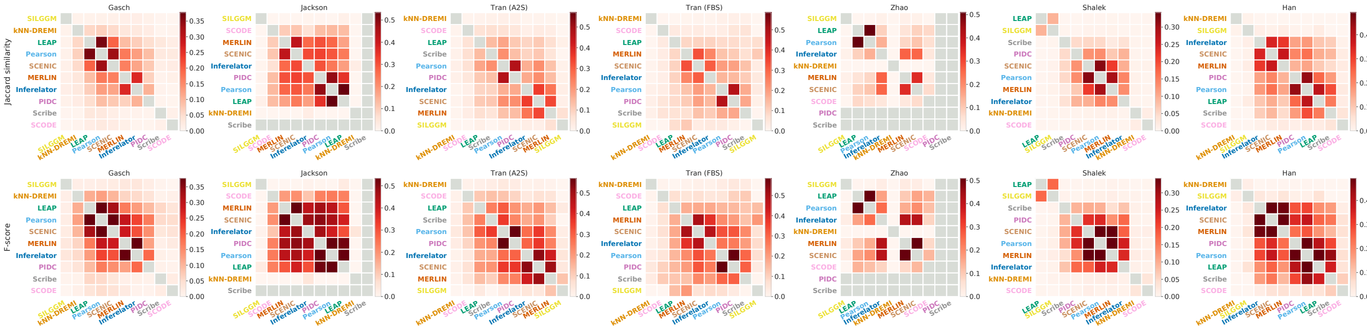
Jaccard similarity
F-score
Gasch
Jackson
Tran (A2S)
Tran (FBS)
Zhao
Shalek
Han

**Figure S10.** Comparison of Jaccard index and F-score as similarity metrics. Top, network similarity matrices using the Jaccard index for top 5k edges, with columns ordered by a hierarchical clustering of the respective matrix. Bottom, similarity matrices using the F-score. The patterns of network similarity are broadly similar when using either metric, while the F-score yields scores of higher magnitude.

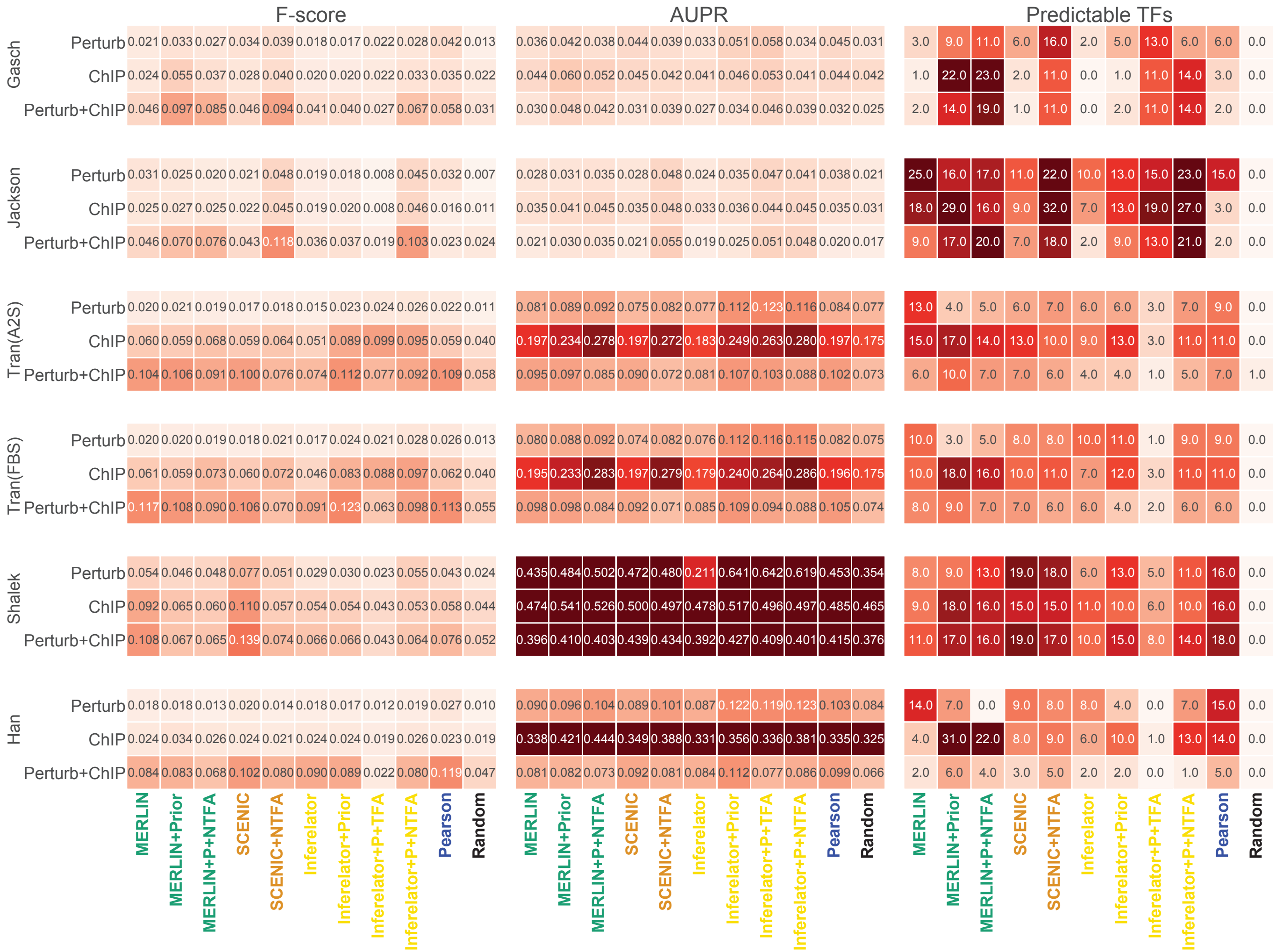
F-score
AUPR
Predictable TFs
Gasch
Jackson
Tran(A2S)
Tran(FBS)
Shalek
Han
Perturb
ChIP
Perturb+ChIP
MERLIN
MERLIN+Prior
MERLIN+P+NTFA
SCENIC
SCENIC+NTFA
Inferelator
Inferelator+Prior
Inferelator+P+TFA
Inferelator+P+NTFA
Pearson
Random
Gasch Perturb 0.021 0.033 0.027 0.034 0.039 0.018 0.017 0.022 0.028 0.042 0.013 | 0.036 0.042 0.038 0.044 0.039 0.033 0.051 0.058 0.034 0.045 0.031 | 3.0 9.0 11.0 6.0 16.0 2.0 5.0 13.0 6.0 6.0 0.0
Gasch ChIP 0.024 0.055 0.037 0.028 0.040 0.020 0.020 0.022 0.033 0.035 0.022 | 0.044 0.060 0.052 0.045 0.042 0.041 0.046 0.053 0.041 0.044 0.042 | 1.0 22.0 23.0 2.0 11.0 0.0 1.0 11.0 14.0 3.0 0.0
Gasch Perturb+ChIP 0.046 0.097 0.085 0.046 0.094 0.041 0.040 0.027 0.067 0.058 0.031 | 0.030 0.048 0.042 0.031 0.039 0.027 0.034 0.046 0.039 0.032 0.025 | 2.0 14.0 19.0 1.0 11.0 0.0 2.0 11.0 14.0 2.0 0.0
Jackson Perturb 0.031 0.025 0.020 0.021 0.048 0.019 0.018 0.008 0.045 0.032 0.007 | 0.028 0.031 0.035 0.028 0.048 0.024 0.035 0.047 0.041 0.038 0.021 | 25.0 16.0 17.0 11.0 22.0 10.0 13.0 15.0 23.0 15.0 0.0
Jackson ChIP 0.025 0.027 0.025 0.022 0.045 0.019 0.020 0.008 0.046 0.016 0.011 | 0.035 0.041 0.045 0.035 0.048 0.033 0.036 0.044 0.045 0.035 0.031 | 18.0 29.0 16.0 9.0 32.0 7.0 13.0 19.0 27.0 3.0 0.0
Jackson Perturb+ChIP 0.046 0.070 0.076 0.043 0.118 0.036 0.037 0.019 0.103 0.023 0.024 | 0.021 0.030 0.035 0.021 0.055 0.019 0.025 0.051 0.048 0.020 0.017 | 9.0 17.0 20.0 7.0 18.0 2.0 9.0 13.0 21.0 2.0 0.0
Tran(A2S) Perturb 0.020 0.021 0.019 0.017 0.018 0.015 0.023 0.024 0.026 0.022 0.011 | 0.081 0.089 0.092 0.075 0.082 0.077 0.112 0.123 0.116 0.084 0.077 | 13.0 4.0 5.0 6.0 7.0 6.0 6.0 3.0 7.0 9.0 0.0
Tran(A2S) ChIP 0.060 0.059 0.068 0.059 0.064 0.051 0.089 0.099 0.095 0.059 0.040 | 0.197 0.234 0.278 0.197 0.272 0.183 0.249 0.263 0.280 0.197 0.175 | 15.0 17.0 14.0 13.0 10.0 9.0 13.0 3.0 11.0 11.0 0.0
Tran(A2S) Perturb+ChIP 0.104 0.106 0.091 0.100 0.076 0.074 0.112 0.077 0.092 0.109 0.058 | 0.095 0.097 0.085 0.090 0.072 0.081 0.107 0.103 0.088 0.102 0.073 | 6.0 10.0 7.0 7.0 6.0 4.0 4.0 1.0 5.0 7.0 1.0
Tran(FBS) Perturb 0.020 0.020 0.019 0.018 0.021 0.017 0.024 0.021 0.028 0.026 0.013 | 0.080 0.088 0.092 0.074 0.082 0.076 0.112 0.116 0.115 0.082 0.075 | 10.0 3.0 5.0 8.0 8.0 10.0 11.0 1.0 9.0 9.0 0.0
Tran(FBS) ChIP 0.061 0.059 0.073 0.060 0.072 0.046 0.083 0.088 0.097 0.062 0.040 | 0.195 0.233 0.283 0.197 0.279 0.179 0.240 0.264 0.286 0.196 0.175 | 10.0 18.0 16.0 10.0 11.0 7.0 12.0 3.0 11.0 11.0 0.0
Tran(FBS) Perturb+ChIP 0.117 0.108 0.090 0.106 0.070 0.091 0.123 0.063 0.098 0.113 0.055 | 0.098 0.098 0.084 0.092 0.071 0.085 0.109 0.094 0.088 0.105 0.074 | 8.0 9.0 7.0 7.0 6.0 6.0 4.0 2.0 6.0 6.0 0.0
Shalek Perturb 0.054 0.046 0.048 0.077 0.051 0.029 0.030 0.023 0.055 0.043 0.024 | 0.435 0.484 0.502 0.472 0.480 0.211 0.641 0.642 0.619 0.453 0.354 | 8.0 9.0 13.0 19.0 18.0 6.0 13.0 5.0 11.0 16.0 0.0
Shalek ChIP 0.092 0.065 0.060 0.110 0.057 0.054 0.054 0.043 0.053 0.058 0.044 | 0.474 0.541 0.526 0.500 0.497 0.478 0.517 0.496 0.497 0.485 0.465 | 9.0 18.0 16.0 15.0 15.0 11.0 10.0 6.0 10.0 16.0 0.0
Shalek Perturb+ChIP 0.108 0.067 0.065 0.139 0.074 0.066 0.066 0.043 0.064 0.076 0.052 | 0.396 0.410 0.403 0.439 0.434 0.392 0.427 0.409 0.401 0.415 0.376 | 11.0 17.0 16.0 19.0 17.0 10.0 15.0 8.0 14.0 18.0 0.0
Han Perturb 0.018 0.018 0.013 0.020 0.014 0.018 0.017 0.012 0.019 0.027 0.010 | 0.090 0.096 0.104 0.089 0.101 0.087 0.122 0.119 0.123 0.103 0.084 | 14.0 7.0 0.0 9.0 8.0 8.0 4.0 0.0 7.0 15.0 0.0
Han ChIP 0.024 0.034 0.026 0.024 0.021 0.024 0.024 0.019 0.026 0.023 0.019 | 0.338 0.421 0.444 0.349 0.388 0.331 0.356 0.336 0.381 0.335 0.325 | 4.0 31.0 22.0 8.0 9.0 6.0 10.0 1.0 13.0 14.0 0.0
Han Perturb+ChIP 0.084 0.083 0.068 0.102 0.080 0.090 0.089 0.022 0.080 0.119 0.047 | 0.081 0.082 0.073 0.092 0.081 0.084 0.112 0.077 0.086 0.099 0.066 | 2.0 6.0 4.0 3.0 5.0 2.0 2.0 0.0 1.0 5.0 0.0

**Figure S11.** Network inference utilizing gene expression alone compared to addition of priors and Transcription factor Activity (TFA). The terms "+P" after an algorithm's name denotes adding prior while addition of "+TFA" or "+NTFA" corresponds to TFA estimated using Inferelator's inbuilt TFA estimation or TFA estimated from Network Components Analysis (NTFA), respectively. The performance of MERLIN, SCENIC, and Inferelator is reported along with their enhanced versions incorporating prior, TFA or both, on 6 datasets. On the left are the values for F-score. The middle column shows values for AUPR. On the right are the number of predictable TFs. For calculating the F-score and predictable TFs we considered only the top 5,000 edges while for AUPR we used all edges. We also report these metrics for Pearson and the Random network.
