## Supplemental Data 3 for "Identifying strengths and weaknesses of methods for computational network inference from single cell RNA-seq data"

**A study on how the size of a dataset impacts the performance of the network inference methods**

**Yeast study.**

For the yeast cell types, we had two datasets – Gasch (163 cells) and Jackson (17,396 cells). We randomly downsampled Jackson to generate 5 subsamples where each subsample contains 163 cells randomly selected from its original 17,396 cells without replacement. Thus, the number of cells in each Jackson subsample exactly matches the number of cells in the Gasch dataset.

| **Dataset** | **Cell type** | **No. of cells** | **No. of genes** |
| --- | --- | --- | --- |
| Gasch | Yeast | 163 | 3,847 |
| Jackson | Yeast | 17,396 | 5,736 |
| Each of the five Jackson subsamples after random downsampling | Yeast | 163 | 5,736 |

We utilized two methods, MERLIN and SCENIC, since they had performed consistently well across different datasets in our previous experiments. These two methods were applied on Gasch, Jackson, and the five distinct subsamples of Jackson. When applying MERLIN on a Jackson subsample, we generated 100 smaller subsamples half the size of the original subsample (i.e. ceiling(163/2) = 82 cells per smaller subsample). We ran MERLIN independently on each of the smaller subsamples, thus inferring 100 networks. We made a consensus network by combining these 100 networks. The consensus network represents MERLIN’s inferred network corresponding to the original subsample. On the other hand, we did not need to do such explicit resampling for SCENIC since it performs bootstrapping internally and produces a consensus network for each of the Jackson subsamples.

We observed that both MERLIN and SCENIC encountered decreased AUPR, F-score, and the number of predictable TFs when the number of cells in Jackson is reduced from 17,396 to 163 (Jackson vs. Jackson subsamples) with only one exception (SCENIC in panel C1 against Jackson subsample 5, **Figure 1**). We also observed that the decrements are lower for SCENIC compared to that of MERLIN. In other words, SCENIC’s performance was less susceptible to the downsampling of the Jackson dataset compared to that of MERLIN.

Subsequently, we compared the performance on the Gasch dataset vs. on the Jackson subsamples since they all have equal number of cells (**Figure 1**). We found that the performance of both MERLIN and SCENIC on the Jackson subsamples are similar to their performance on the full Jackson dataset in relation to the Gasch dataset. For example, with respect to the Perturb gold standard network, SCENIC has a lower AUPR score for the full Jackson dataset compared to that for the Gasch dataset; similarly it has lower AUPR scores for the Jackson subsamples compared to that for the Gasch dataset. On the other hand, SCENIC has a higher number of predictable TFs for the full Jackson dataset compared to that for the Gasch dataset; similarly again, it has higher numbers of predictable TFs for the Jackson subsamples compared to that for the Gasch dataset. Therefore, the performance of a method is not necessarily similar on two distinct datasets simply because the said datasets have the same number of cells.

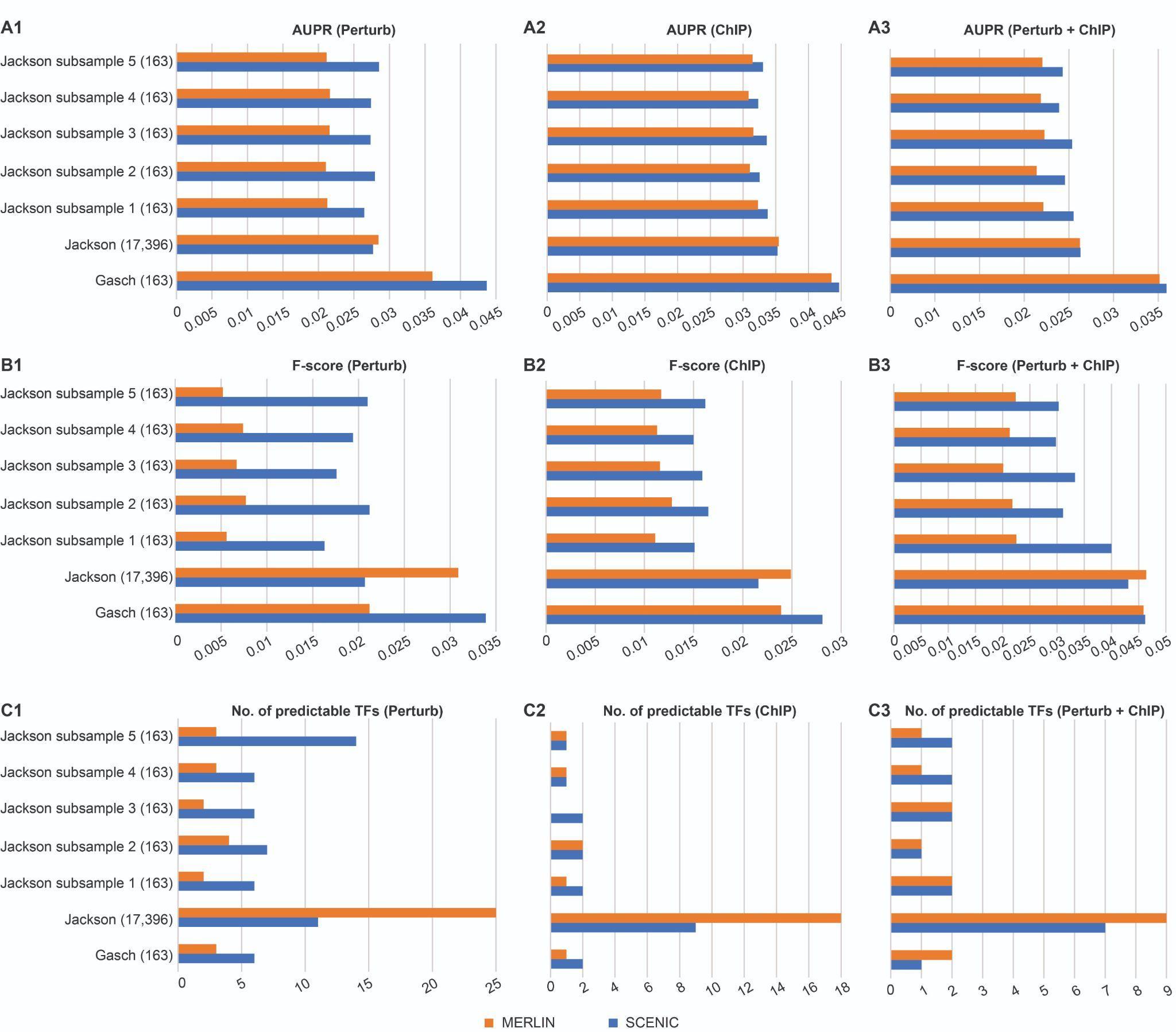

**Figure 1**. The performance of MERLIN and SCENIC on Gasch, Jackson, and the five different subsamples of the Jackson dataset with respect to three different gold standard networks – Perturb, ChIP, and their common edges (Perturb + ChIP). The number of cells in a dataset is mentioned in the parentheses by its name. For calculating the F-score and the number of predictable TFs of an inferred network, we ranked its edges by their predicted confidence scores and considered only the top 5,000 edges.

**mESC study.**

For the mammalian cell types, we utilized the mESC datasets namely the Tran (FBS) dataset (3,324 cells) and Zhao dataset (36,199 cells). We randomly downsampled Zhao to generate 5 subsamples where each subsample contains 3,324 cells randomly selected from its original 36,199 cells without replacement.

| **Dataset** | **Cell type** | **No. of cells** | **No. of genes** |
| --- | --- | --- | --- |
| Tran (FBS) | mESC | 3,324 | 6,621 |
| Zhao | mESC | 36,199 | 8,442 |
| Each of the five Zhao subsamples after random downsampling | mESC | 3,324 | 8,442 |

We followed the same network inference pipelines with MERLIN and SCENIC as done in the yeast study. The results are presented in **Figure2**. We observed that, with respect to the Perturb and Perturb+ChIP gold standard networks, both MERLIN and SCENIC encountered decreased AUPR when the number of cells in Zhao is reduced from 36,199 to 3,324 (Zhao vs. Zhao subsamples). However, with respect to the ChIP gold standard network, their AUPR scores increased. The F-scores and numbers of predictable TFs generally decreased after Zhao downsampling. This is possible if the true positive edges of ChIP gold standard network are not among the top 5,000 edges in the inferred networks (since F-score and the number of predictable TFs are calculated on only the top 5,000 edges unlike AUPR which is calculated on the whole inferred network). Therefore, there does not exist a consistent pattern of increment or decrement across different metrics that can be attributed to the downsampling of the Zhao dataset.
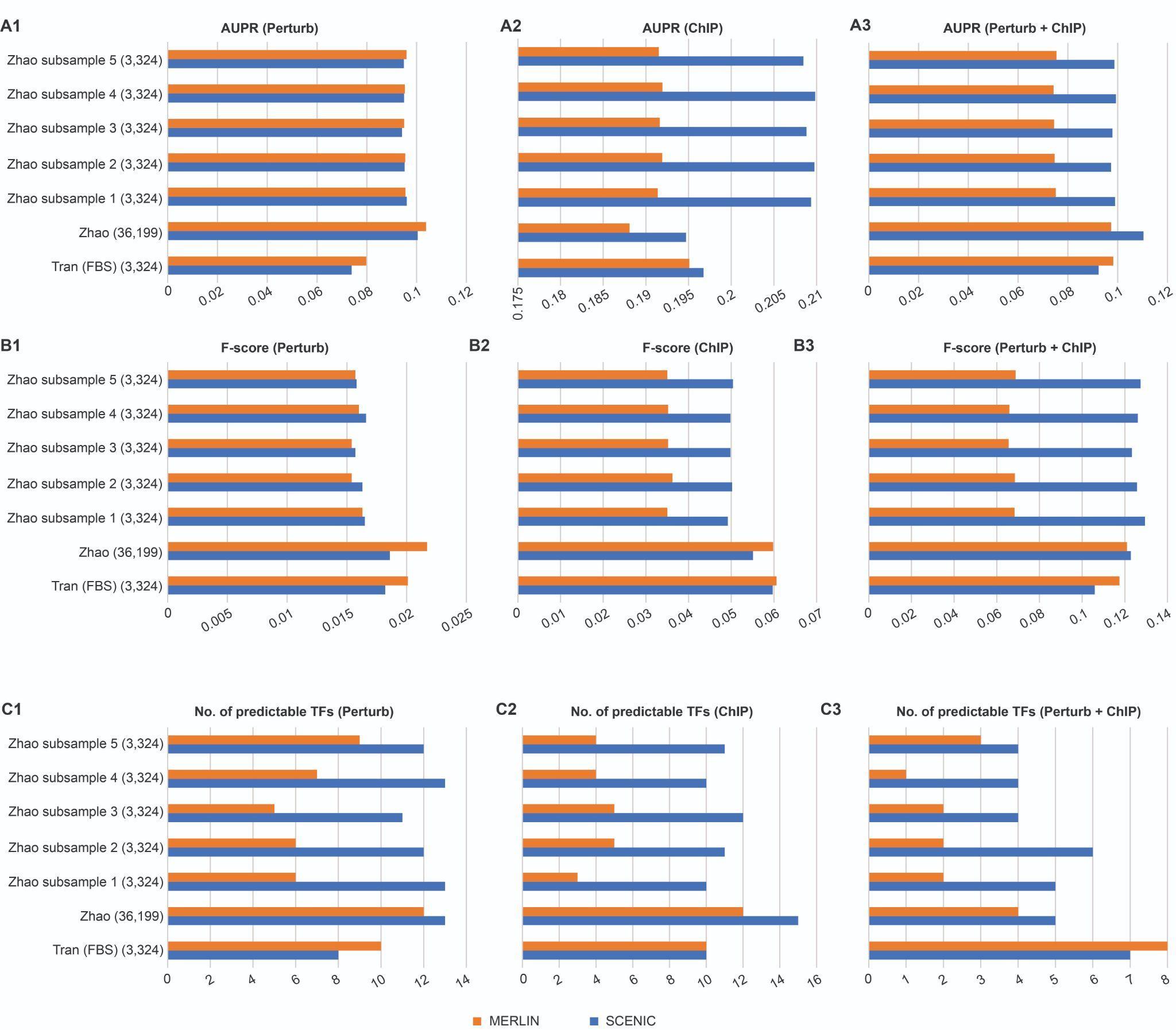
**Figure 2.** The performance of MERLIN and SCENIC on Tran (FBS), Zhao, and the five different subsamples of the Zhao dataset with respect to three different gold standard networks – Perturb, ChIP, and their common edges (Perturb + ChIP). The number of cells in a dataset is mentioned in the parentheses by its name. For calculating the F-score and the number of predictable TFs of an inferred network, we ranked its edges by their predicted confidence scores and considered only the top 5,000 edges.

From the above observations, we made the following conclusions:

- The performance of a method was not necessarily similar on two distinct datasets simply because the datasets have the same number of cells, e.g., Gasch vs. Jackson subsamples, Tran (FBS) vs. Zhao subsamples. One of the factors could be the difference in the sequencing depths of the datasets. We observed that the total number of transcripts (across all genes) per cell and the number of distinct genes detected per cell are higher in Gasch compared to that of the Jackson subsamples (**Figure 3**, panels A1-A2). It supports the observation that both SCENIC and MERLIN consistently achieved higher AUPR and F-scores with Gasch compared to that with the Jackson subsamples. On the other hand, for Tran (FBS) vs. the Zhao subsamples, although the total number of transcripts per cell is higher in Tran (FBS), the number of distinct genes detected per cell is higher in the Zhao subsamples (**Figure 3**, Panels B1-B2). It adds support to the observation that none of the methods consistently perform better for either Tran (FBS) or the Zhao subsamples.
- Even when a method is applied on a single dataset and its downsized subsamples, the effect of downsampling did not have a consistent positive or negative effect on the evaluation metrics. For Jackson, downsampling from 17,396 cells to 163 cells had a negative effect (less so for SCENIC than MERLIN), whereas, for Zhao, downsampling from 36,199 cells to 3,324 cells did not have any consistent effect. The number of cells in the downsized subsample might also be a factor. For Zhao, even the downsized subsamples have 3,324 cells which is a significantly higher number compared to only 163 cells in the downsized subsamples of Jackson. Moreover, for Zhao, we observed that higher AUPR scores can be achieved with downsized subsamples compared to that with the full dataset. We cannot rule out greater noise in the Zhao dataset and the mammalian gold standards which can contribute to these patterns. We note that SCENIC was less susceptible to downsampling compared to MERLIN which suggests the tree-based non-parametric modeling of SCENIC is beneficial for robust performance on scRNA-seq datasets which tend to be noisy.
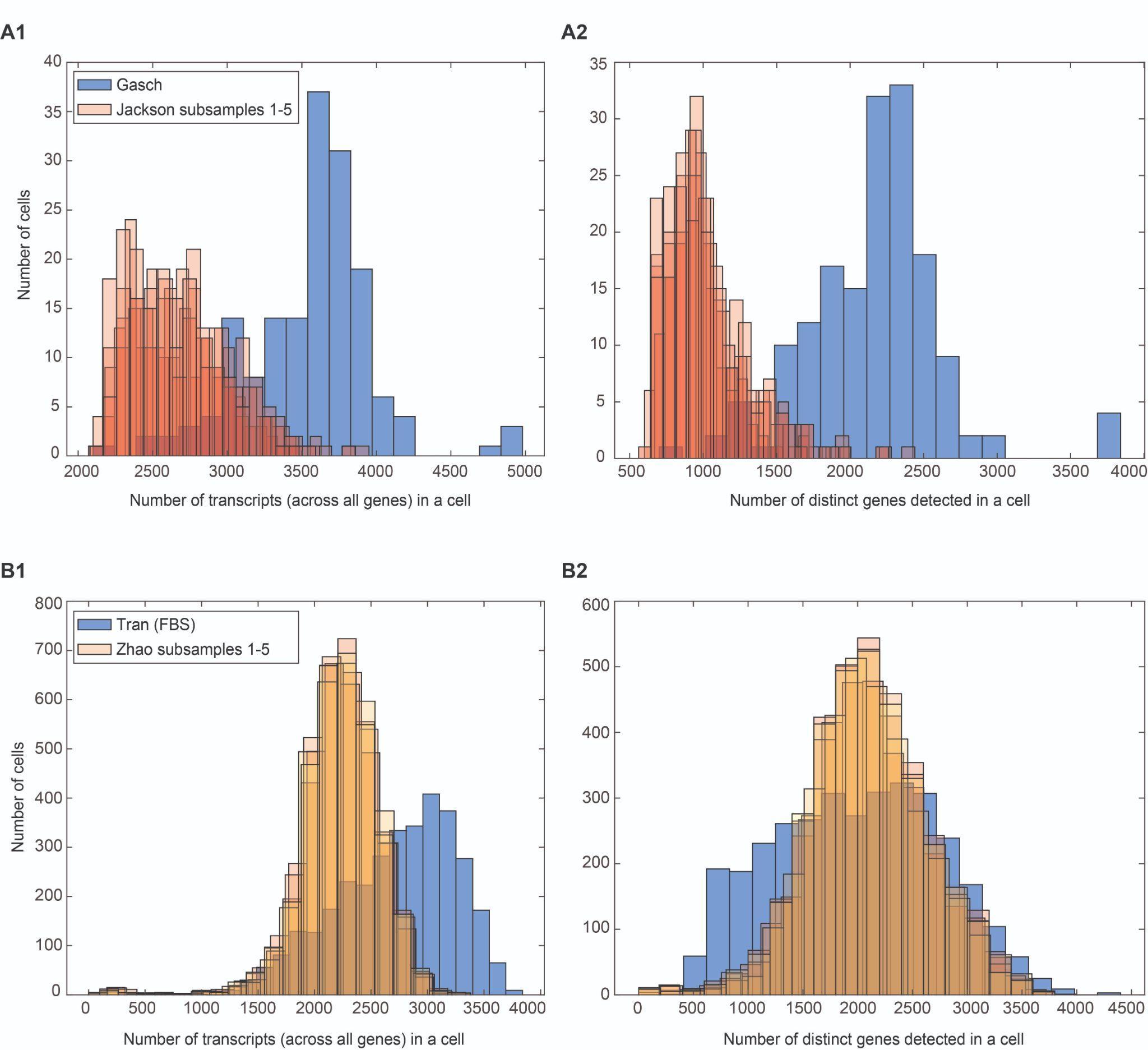

**Figure 3.** The comparison of sequencing depths between datasets of the same size (the same number of cells). (A1-A2) presents the comparison between Gasch and five Jackson subsamples. Each of the Jackson subsamples has 163 cells which is the same number of cells Gasch has. However, the Jackson subsamples differ from the Gasch dataset in terms of (A1) the total number of transcripts (read across all genes) in each cell and (A2) the number of distinct genes detected in each cell. Similarly, (B1-B2) presents the differences in the total number of transcripts (across all genes) in each cell as well as the number of distinct genes detected in each cell of Tran (FBS) and the Zhao subsamples, even though each of the datasets has the same number of cells (3,324 cells).
